# Single-Cell Profiling of the Developing Organ of Corti Identifies Etv4/5/1 as Key Regulators of Pillar Cell Identity

**DOI:** 10.64898/2026.01.19.700450

**Authors:** Susumu Sakamoto, Matthew W. Kelley

## Abstract

The mammalian auditory sensory epithelium, the organ of Corti, contains a number of unique cell types, including the inner and outer pillar cells that form the walls of the tunnel of Corti. The limited number of pillar cells and their close physical proximity to other cells within the organ of Corti has limited efforts to transcriptionally characterize their development. To identify potential regulators of pillar cell formation, we isolated cochlear duct epithelial cells at time points between embryonic day 11 and 16 for single cell RNA-sequencing. The resulting data were used to build a developmental trajectory from undifferentiated precursor cells to each of the major cell types in the organ of Corti including inner pillar cells. Bioinformatic analyses including SCENIC, TradeSeq and CellOracle identified the Etv4/5/1 transcription factors as likely regulators of inner pillar cell development. To specifically examine the role of Etv4/5/1 in pillar cell development, conditional triple mutants were generated. Results indicate defects in the formation of inner pillar cells as well as changes in other cochlear cells likely as a result of secondary interactions. Finally, we demonstrate that expression of *Etv4/5/1* in pillar cells is dependent on *Fgfr3* and identify downstream targets of Fgfr3/Etv signaling in inner pillar cells. These results provide significant insights regarding the specification and early development of inner pillar cells, which will have implications for understanding congenital deficits and potential applications in the development of regenerative strategies.

## Introduction

The mammalian auditory sensory epithelium, the organ of Corti (OC), is a narrow strip of mechanosensory hair cells (HCs) and associated supporting cells (SCs) extending along the length of the cochlear spiral. At the cellular level, the OC is asymmetrically organized along it’s medial-to-lateral axis^1–3^. The medial side of the OC contains a single row of IHCs which are surrounded by inner phalangeal cells (IPhCs) and border cells (BCs), while the lateral side of the OC is comprised of three rows of outer hair cells (OHCs) which are surrounded by Deiters’ cells (DCs). The medial and lateral regions are separated by single rows of IPCs and OPCs which together form the tunnel of Corti^1,2,4,5^ While IHCs, IPhCs and BCs are phenotypically similar to HCs and SCs located in other inner ear sensory epithelia, the OHCs, DCs, IPCs and OPCs represent unique cell types that are only found in the OC. The evolution of these cell types occurred concurrently with an increase in frequency range and acuity in the mammalian lineage^6^. While recent studies have identified transcription factors that play a role in development of OHCs^7,8^, the genetic regulation of the development of other OC cell types, such as the pillar cells, has not been determined.

The cells that will develop as pillar cells and all other cells within the OC are derived from a population of precursor cells, termed prosensory cells (PsCs), located in the middle region of the cochlear duct, an epithelial structure that extends from the ventral region of the otocyst beginning around E11 in the mouse ^9,10^. At early developmental time points, the cells in the floor of the duct appear homogenous, however gene expression and fate-mapping studies have demonstrated differences along the medial-to-lateral axis of the spiral prior to morphological differentiation (Fig. 1a)^10,11^. In particular, the HMG-box transcription factor SOX2 is expressed across the medial two-thirds of the duct at E11 and E12 before becoming restricted to a band of cells located in the middle of the duct ^12–14^. All the cell types located within the OC are believed to arise from this population of SOX2^+^-prosensory cells ^10^. Moreover, recent studies have demonstrated that OHCs, IPCs, OPCs, and DCs arise from a spatially distinct population of progenitors termed lateral prosensory cells (Lat.PsCs) that become specified around E14^15^. While Lat.PsCs show a strong bias towards cell fates located in the lateral compartment of the OC, recent results have demonstrated that changes in expression of the transcription factors TBX2 or INSM1 can lead to phenotypic transformations in which OHCs convert into IHCs or vice versa ^7,16,17^. These results indicate that while cell fates within the developing OC are not immutable, unique OC cell types normally arise from medial or lateral prosensory cells, suggesting that individual precursors are biased towards specific cell fates, a conclusion that is supported by recent lineage tracing studies^11^. Several recent studies have examined the factors that might play a role in specifying different cellular domains along the medial-to-lateral axis of the duct ^18–20^, but the developmental origins of individual cell types, and in particular pillar cells, have not been determined. Similarly, while scRNAseq datasets for different time points in cochlear development have recently been published^21–23^ and made available on gEAR (umgear.org), the time period between E11 and E16 had not been analyzed specifically. Therefore, in this study we generated a single cell RNA sequencing (scRNAseq) dataset focused on the developmental progression of cochlear prosensory cells to their final cellular phenotypes. Based on the results of that analysis, the Etv4/5/1 transcription factors were identified as likely regulators of IPC fates. To test that hypothesis, we generated conditional *Etv4/5/1* triple knock out mice. Results demonstrated a loss of IPCs in the absence of Etv signaling with downstream changes in development of other OC cell types including OHCs and OPCs. Finally, the expression of *Etv4/5/1* was shown to be dependent on Fgfr3 and early targets of Fgfr3/Etv signaling were identified.

**Figure 1.**
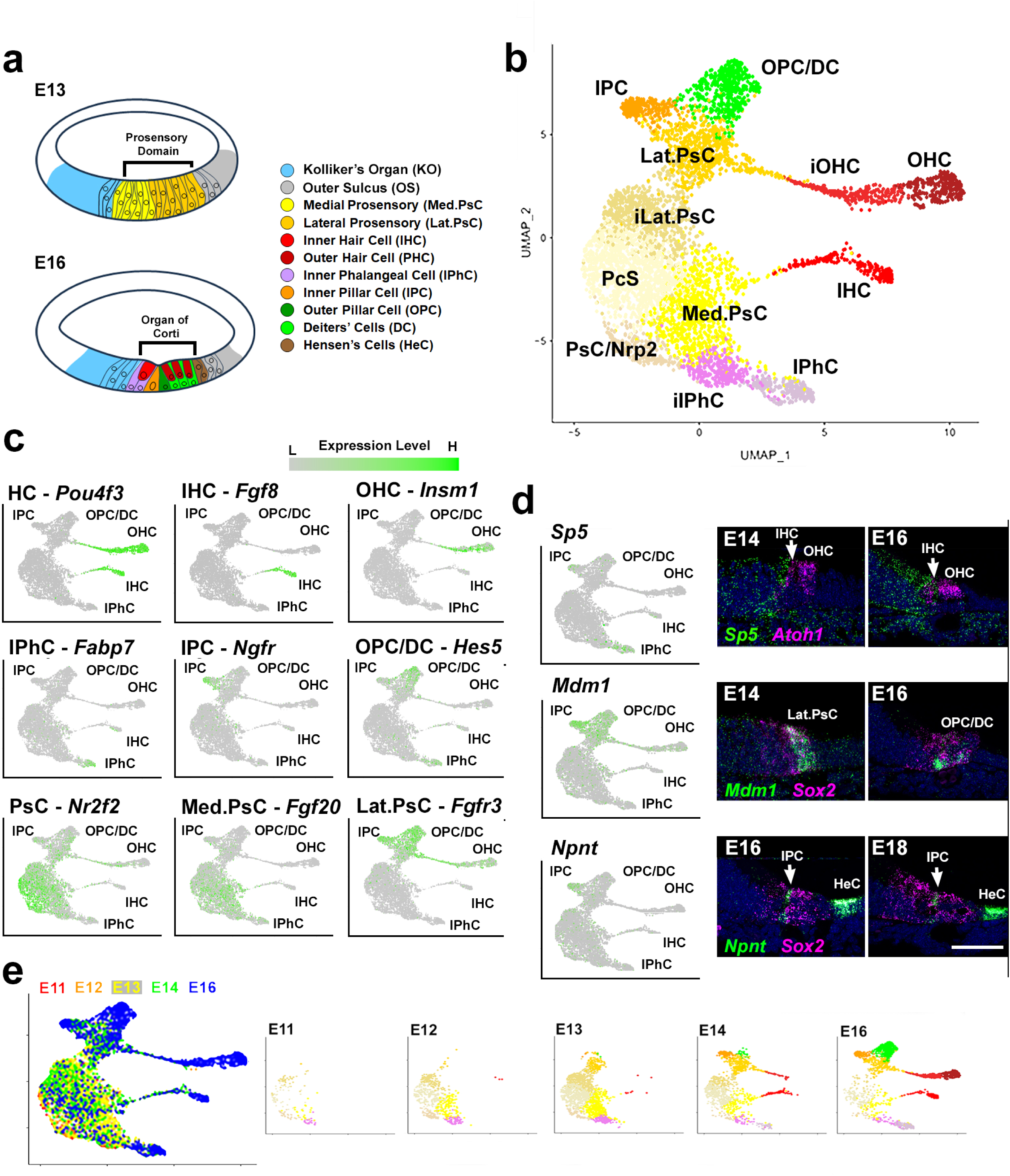
Developmental of the OC. **a.** Line drawings illustrating the structure of the cochlear duct at E13 and E16. Cell types are indicated. See text for details. **b.** UMAP projection of ∼5500 single cells isolated from the developing OC between E11 and E16. Known cellular phenotypes, including PsCs, IHCs, OHCs, IPhCs, IPCs and OHCs/DCs are indicated. Other clusters appear to represent transitional progenitor states. **c.** Feature plots for marker genes for each of the known cell clusters in b**. d.** Feature plots and smFISH in cryosections for three novel markers of different cochlear cell types. *Sp5* is expressed in IPhCs, *Mdm1* is expressed in IPCs, OPC/DCs and OHCs. *Npnt* is expressed in IPCs and in HeCs which were not included in the OC sensory dataset. **e.** Left panel shows the same plot as in b but labeled based on day of collection. Remaining panels show cell types collected at each developmental time point (color scheme as in panel b). Abbr. Med.PsC: medial prosensory cells, iIPhC: intermediate inner phalangeal cells, iOHC: intermediate OHCs, Lat.PsC; intermediate lateral prosensory cells, Lat.PsC: lateral prosensory cells. Scale bar in d (same for all smFISH images), 50 μm.

## Results

### Characterization of the cellular population in the early cochlear duct

Single cells were isolated and captured from the cochlear ducts of CD1 embryos at embryonic days 11 (E11), 12, 13, 14 or 16. Following quality control analysis to eliminate doublets and cells under stress, a total of just over 43,000 cells (4,362 at E11, 7,746 at E12, 9,143 at E13, 8,825 at E14, 13,573 at E16) were included in the data set. Next, differential gene expression was used to generate a subset of the data containing only PsCs and OC cell types. An analysis of the full dataset will be provided in a subsequent publication. PsCs and OC cells from each time point (just over 5,500 cells total, mean number of genes per cell = 2,607) were then combined to form a single OC Sensory Dataset (Fig. 1b and Data File 1). Unbiased clustering identified twelve transcriptionally distinct clusters. Subsequent analysis of gene expression for known cell types identified clusters corresponding to specific OC cell types including IHCs, OHCs, OPCs/DCs, IPCs and IPhCs as well as PsCs, Med.PsCs and Lat.PsCs^7,24–31^ (Fig. 1b,c, Suppl. Fig. 1, and Data File 2). BCs were not identified as a unique cluster and were assumed to co-cluster with IPhCs as these cells are likely to be very similar transcriptionally. Similarly, in contrast with scRNAseq data from P1 cochleae^22,32^, at this stage OPCs and DCs formed a single cluster, which may reflect transcriptional similarity prior to E16^22,32^, or limitations in the cellular resolution of our method of analysis.

Next, to evaluate the quality of the dataset, expression of three genes that had not been previously examined in the OC were localized using single molecule fluorescent in situ hybridization (smFISH). Feature plots for *Sp5, Mdm1* and *Npnt* indicated restricted expression to IPhCs, all lateral cell types and IPCs, respectively (Fig. 1d). smFISH for each gene in cochlear cross sections from E14 and E16 confirmed the feature plot results. In addition to expression in IPCs, *Npnt* was also observed in developing Hensen’s cells (HeCs) located lateral to the OC.

### Developmental Trajectory Analysis

The relative positions of the identified cell clusters on the UMAP projection suggested a potential developmental trajectory that bifurcated from the PsC cluster to give rise to Lat.PsC and Med.PsC clusters which then give rise to the known OC cell types. To characterize this possible trajectory, the UMAP was replotted based on day of collection (Fig. 1e). Results indicated that PsC cell types were the predominant cell types collected between E11 and E13. In contrast, HC and SC cell types were almost exclusively collected at E14 or E16. Next, we used UniTVelo, a modification of RNA velocity^33^, to determine the overall flow of cellular development. Results indicated a developmental flow consistent with the analysis of cell type composition based on age of collection (Fig. 2a). PsCs flow toward the Lat.PsC and Med.PsC clusters which then give rise to known OC cell types. Based on these results, we used Slingshot ^34^ to identify potential developmental trajectories within the dataset. Consistent with the previous analyses, the Slingshot results indicated an initial population of PsCs that splits into two trajectories (Fig. 2b). A medial trajectory that gives rise to Med.PsCs which then split into IHCs and IPhCs and a lateral trajectory that gives rise to Lat.PsCs prior to developing into OHC, IPC, and OPC/DC clusters (Fig. 2b).

**Figure 2.**
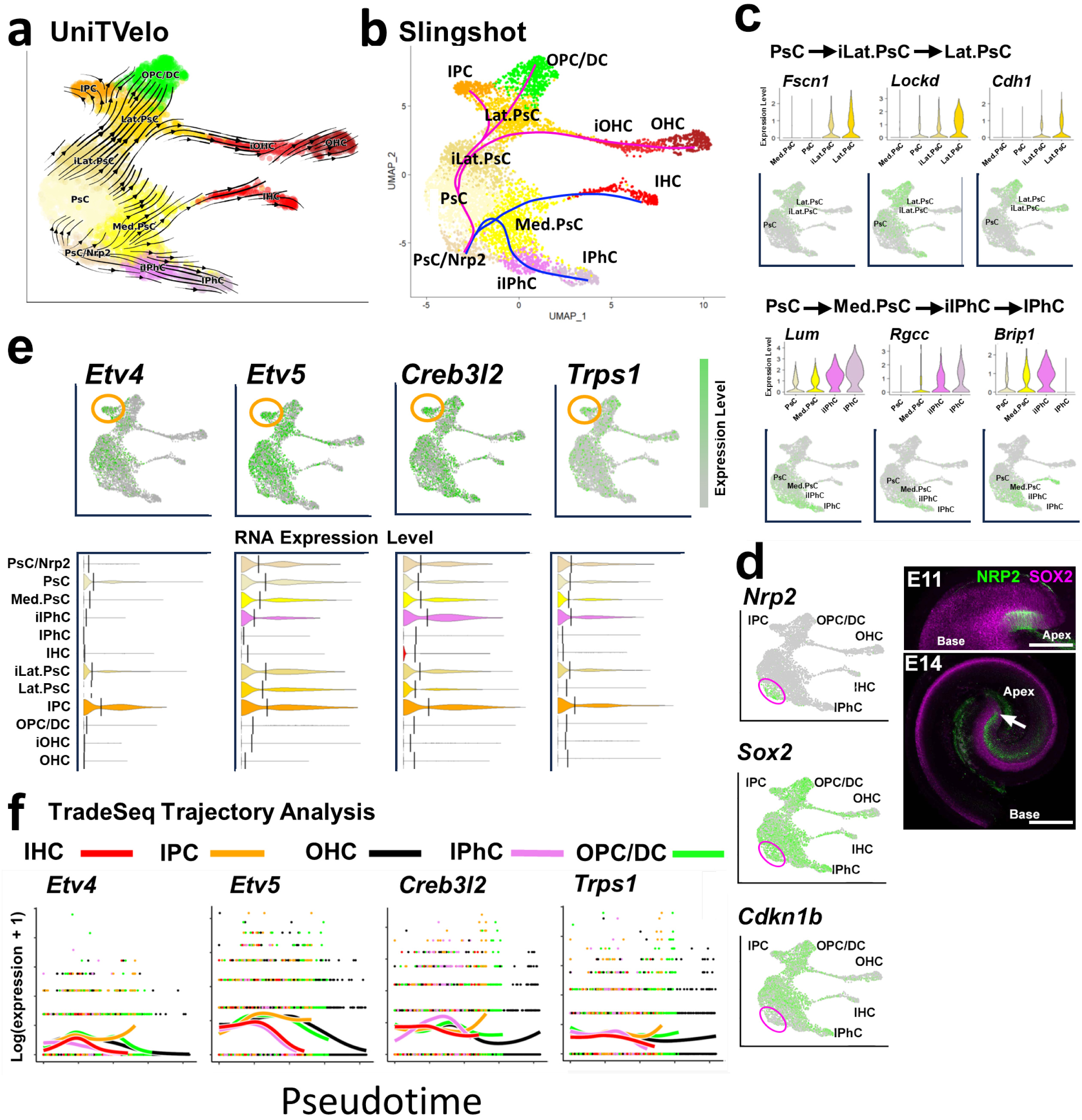
Developmental Trajectory for OC cell types. **a.** UniTVelo trajectory analysis. Cluster identities are as in Fig. 1a. Arrows indicate developmental flow. Overall flow is from the PsC cluster along a lateral trajectory to the Lat.PsC and a medial trajectory to the Med.PsC. Psc/Nrp2 cluster flows to both the PsC cluster and to the iIPhC cluster. **b.** Slingshot-generated trajectory indicates similar trajectories to UniTVelo, a medial trajectory (blue) giving rise to IHCs and IPhCs and a lateral trajectory (magenta) giving rise to OHCs, IPCs and OPCs/DCs. The lateral trajectory appears to pass through two transitional phases, iLat.PsC and Lat.PsC (see Fig. 1). **c.** Violin plots and feature plots illustrating developmental progression in gene expression through the iLat.PsC and iIPhC clusters. Graded expression of each gene is consistent with the location of the iLat.PsC and iIPhC clusters as transitional cell types. **d.** Left column: feature plots for *Nrp2, Sox2* and *Cdkn1b*. The *Nrp2* cluster, circled, is positive for *Sox2* but negative for *Cdkn1b*. Right column: whole-mount images of the cochlea at E11 and E14. At E11 NRP2^+^-cells are located in the medial/apical region of the cochlea. Most appear to be SOX2^+^. At E14, NRP2^+^-cells extend towards the base of the cochlea. Most of these cells are negative for SOX2 but a small group at the apex appear to be positive for both NRP2 and SOX2 (arrow). **e.** Feature and violin plots for four transcription factors that were found to have specific or elevated expression in the IPC cluster (orange circle). Lines on violin plots indicate mean value. **f.** TradeSeq analysis for changes in gene expression for the same four genes as in d along the five different developmental trajectories. *Etv4* and *Creb3l2* show a specific increase in expression in IPCs. Scale bar in d, 200 μm.

In addition to the cell clusters described in the previous section, three additional clusters were identified (Fig. 1b). Based on their position on the UMAP and the results of the Slingshot and UniTVelo analysis, these clusters were initially designated as intermediate Lateral Prosensory Cells (iLat.PsC), intermediate Inner Phalangeal Cells (iIPhC), and Prosensory Cells expressing Nrp2 (PsC/Nrp2). To characterize the iLat.PsC and iIPhC clusters, both of which appear to represent transitional cell types, DE gene expression was examined across the PsC, iLat.PsC, Lat.PsC and PsC, Med.PsC, iIPhC, IPhC trajectories. Consistent with the position of these clusters on the UMAP, results indicated multiple genes showing graded changes in expression across both trajectories (Fig. 2c and Suppl. Fig. 2).

Cells from the final novel cluster, the PsC/Nrp2 cluster, were collected as early as E11 and were located at the beginning of the developmental trajectory in both the UniTVelo and Slingshot analyses. Cells in this cluster showed significant overlap in gene expression with PsCs, including *Sox2*, although not *Cdkn1b*^35^. These cells also expressed a number of other unique genes including *Cyp26a1, Kcnip4* and *Cnr1* (Fig. 2d and Suppl. Fig 3). To determine the spatial location of these cells, expression of NRP2 was localized in the cochlear duct at E11 and E14 (Fig. 2d). Results indicate expression of NRP2 in a small group of cells located at the extreme apex of the cochlea on the medial side at E11. Most of these cells are positive for SOX2. At E14, NRP2^+^-cells extend from the apex of the cochlea towards the base. Most of these cells appear to be located in Kolliker’s organ but a few cells at the extreme apex appear to still be double positive for SOX2. Understanding the relevance of this cell population will require additional characterization. Overall, the initial analysis of this dataset is consistent with the current understanding of cochlear development in which a common pool of PsCs splits into unique Medial and Lateral PsC populations which then give rise to specific subsets of cochlear cell types.

### Development of inner pillar cells

Having characterized the clusters and developmental trajectories within the OC Sensory Dataset, we next focused on the development of IPCs. As discussed, pillar cells are evolutionarily novel cell types that are crucial for the function of the OC. Since the results of the UniTVelo and Slingshot analyses identified a unique trajectory for the development of IPCs, we wanted to use these data to identify factors that might regulate IPC development. As a first step, we identified a total of 24 transcription factors in the top 500 DE genes in the IPC cluster (Data File 2). Of these 24 transcription factors, only four, *Etv4, Etv5*, *Creb3l2* and *Trps1*, showed increased or specific expression in the IPC cell cluster (Fig. 2e and Suppl. Fig. 4). To determine if any of these transcription factors showed a specific increase along the IPC trajectory, we examined expression of all 24 using TradeSeq (Fig. 2e and Suppl. Fig. 5). Of the 24 transcription factors, only *Etv4* and *Creb3l2* showed specific increases along the IPC trajectory (Fig. 2f).

### SCENIC analysis of transcriptional regulons

Because expression of mRNA for a transcription factor is not a definitive indicator of transcriptional activity, we next used SCENIC^36^ to identify expression of transcriptional regulons in each cell cluster (Fig. 3a, Suppl. Fig. 6 and Data File 3). The top ten regulons (based on RSS score) expressed in the IPC cluster were then compared with the top ten regulons expressed in each of the other mature cell type clusters to identify regulons that were likely to be active only in the IPC cluster. Results identified five regulons, Etv4, Etv5, Nkx6-1, Zfp410 and Zfp420, as uniquely active in the IPC cluster (Fig. 3a). Activation of the Creb3l2 regulon was not identified in the IPC cluster and was therefore not pursued further. Feature plots reflecting overall activation of each regulon indicated activation of the Etv4 and Etv5 regulons in the IPC cluster (Fig. 3b). In contrast, feature plots for the Nkx6-1, Zfp410 and Zfp420 regulons indicated minimal activation of these regulons in the IPC cluster (Fig. 3c). To confirm that Nkx6-1, Zfp410 and Zfp420 transcriptional regulons were not specifically active in IPCs, expression of target genes for each those regulons was examined. Results indicated minimal expression of any of the target genes for Nkx6.1, Zfp410 or Zfp420 in the IPC cluster (not shown).

**Figure 3.**
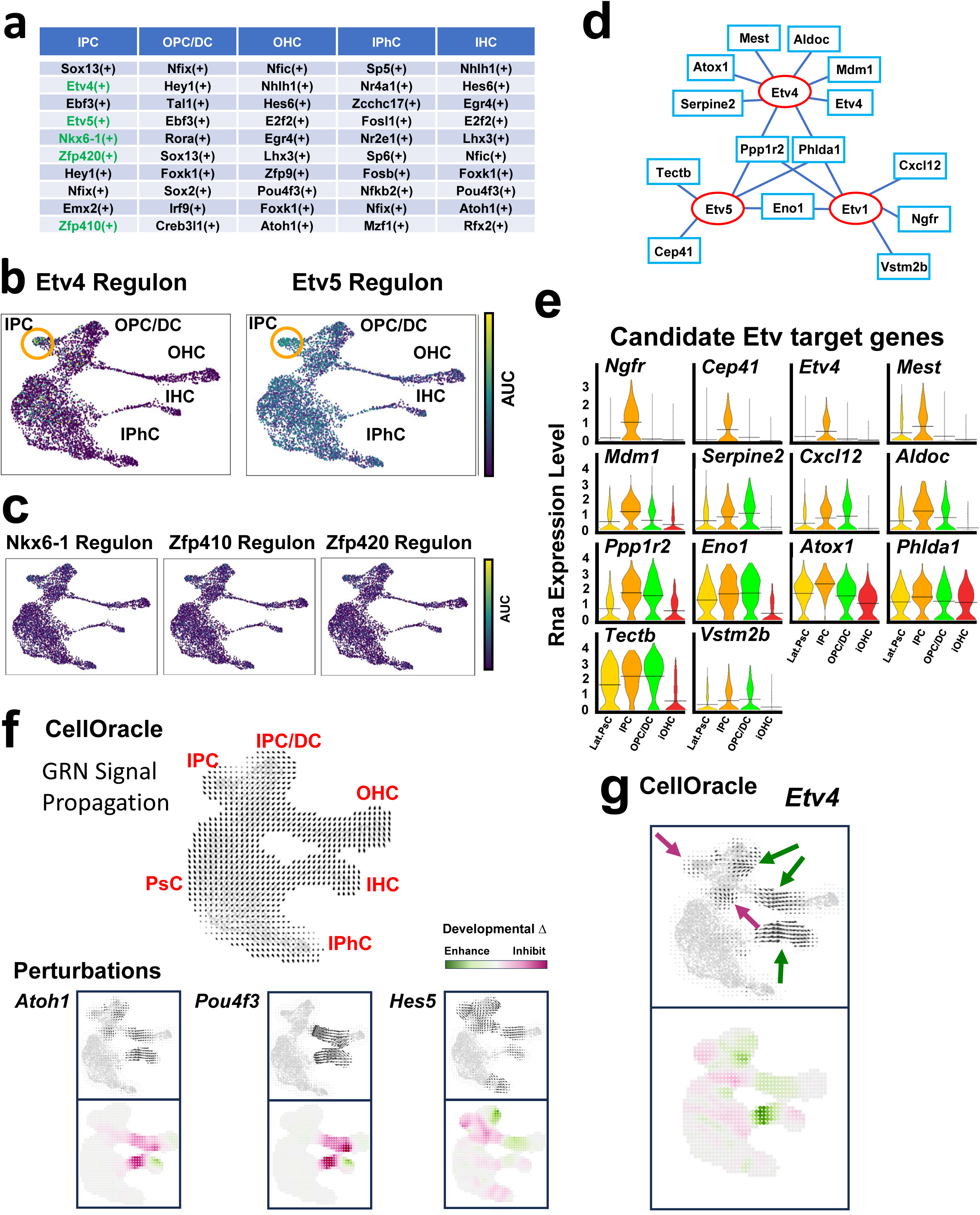
Etv regulons are expressed and likely active in IPCs. **a.** List of top ten regulons (based on RSS) from SCENIC+ analysis of OC development. IPCs express five regulons (in green) that were not among the top ten regulons for any other cell type. **b.** Feature plots showing AUC values for Etv4 and Etv5 regulons. Both are increased in the IPC cluster (orange circles). **c.** Feature plots showing AUC values for Nkx6-1, Zfp410 and Zfp420. None show obvious elevation in the IPC cluster. **d.** Etv gene network developed by combining the list of target genes in the Etv regulons with the DE gene list for IPCs. **e.** Violin plots for the 14 genes listed in d. Four are IPC-specific (orange). **f.** CellOracle analysis. Upper panel: GRN signal propagation arrows indicated developmental flow along the medial and lateral trajectories. Perturbations: Upper panels, arrows indicate predicted changes in developmental trajectories for three genes known to be involved in cochlear development, *Atoh1, Pou4f3, Hes5*. Lower panels, subtraction of perturbed vector flow from GRN signal propagation indicates regions of developmental enhancement (green) or inhibition (magenta). **g.** CellOracle analysis for perturbation of *Etv4*. Results indicate inhibition of IPC and Lat.PsC development and possible enhancement of OPC/DC and OHC development.

Based on the results described above and previous work demonstrating expression of *Etv4* and *Etv5* in developing pillar cells^37,38^, we focused on Etv4 and Etv5 as potential regulators of IPC development. As a first step, we compared the DE gene list for the IPC cluster (Data File 2) with genes within the Etv4/5/1 (because of potential functional redundancy) SCENIC regulons^36^ to identify a list of likely Etv target genes in IPCs (Fig. 3d). Violin plots for each of the top candidate Etv-target genes were generated for the IPC, Lat.PsC, OPC/DC and iOHC clusters (Fig. 3e). Results indicate either exclusive or increased expression of all 14 putative Etv-targets in the IPC cluster (Fig. 3e).

### CellOracle analysis of transcriptional activation and developmental perturbations

The recently introduced CellOracle analysis package incorporates both scRNAseq and, if available, ATACseq, data to identify potential positive and negative transcriptional regulators of cellular development^39,40^. Moreover, CellOracle can also predict changes in a developmental trajectory in response to simulated deletion of a transcription factor. To determine whether CellOracle might be useful in predicting the effects of deletion of *Etv* genes, we first tested the accuracy of this application by simulating deletions of three transcription factors, Atoh*1*, Pou*4f3* or *Hes5,* with known roles in cochlear development. Consistent with previous publications^24,28,41,42^, the main CellOracle predictions for deletions of *Atoh1* or *Pou4f3,* perturbations of hair cell development, were correct (Fig. 3f). However, for both *Atoh1* and *Pou4f3*, limited enhancements of IHC development, which have not been demonstrated in cochlear tissue, were also predicted. For deletion of *Hes5*, CellOracle accurately predicted significant defects in development of SCs but also predicted enhancement of OPC/DC and OHC development. These results also do not reflect known effects of Hes5 deletion (Fig. 3f) ^28,42^.

Since the results of the CellOracle perturbation analysis using previously published cochlear genes predicted both known and either unknown, or possibly incorrect, downstream effects, we wanted to further examine the accuracy of this application by modeling deletion of *Etv4* (Fig. 3g). Results predicted delayed development of IPCs (magenta arrows in Fig. 3g) and enhanced development of OPCs/DCs, IHCs, and OHCs (green arrows in Fig. 3g).

### *Etv4, Etv5* and *Etv1* are expressed in the developing cochlea

To explore the potential role of Etv transcription factors in IPC development, smFISH was performed for *Etv4, Etv5* and *Etv1* in whole mounts and cryosections of the developing cochlea at E14, E16 and E18 (Fig. 4a,b and Suppl. Fig. 7). *Sox2* was used to localize the prosensory domain^12^. Because the OC develops in a basal-to-apical gradient^43^, images were generated for basal, and apical cochlear turns. At E14, low magnification images of whole cochleae indicate broad expression of all three *Etv* genes along the length of the cochlear spiral. At higher magnification, the Etv expression patterns in the more mature basal region of the cochlea clearly include cells expressing *Sox2*, indicating expression in the prosensory domain as well as in the lateral outer sulcus (OS) (Fig. 4a, center column). In the less mature apical region, the PS and OC domains of Etv expression are essentially merged to create a single band of expression encompassing both domains. Cryosections through the cochlear duct confirm the surface images, demonstrating broad expression of each *Etv* gene in the less mature apical region and more restricted expression in the more mature base. As was observed in surface images, in basal sections each *Etv* gene appears to be expressed in a subset of *Sox2*^+^cells and laterally in the non-sensory OS region (Fig. 4a, left column).

**Figure 4.**
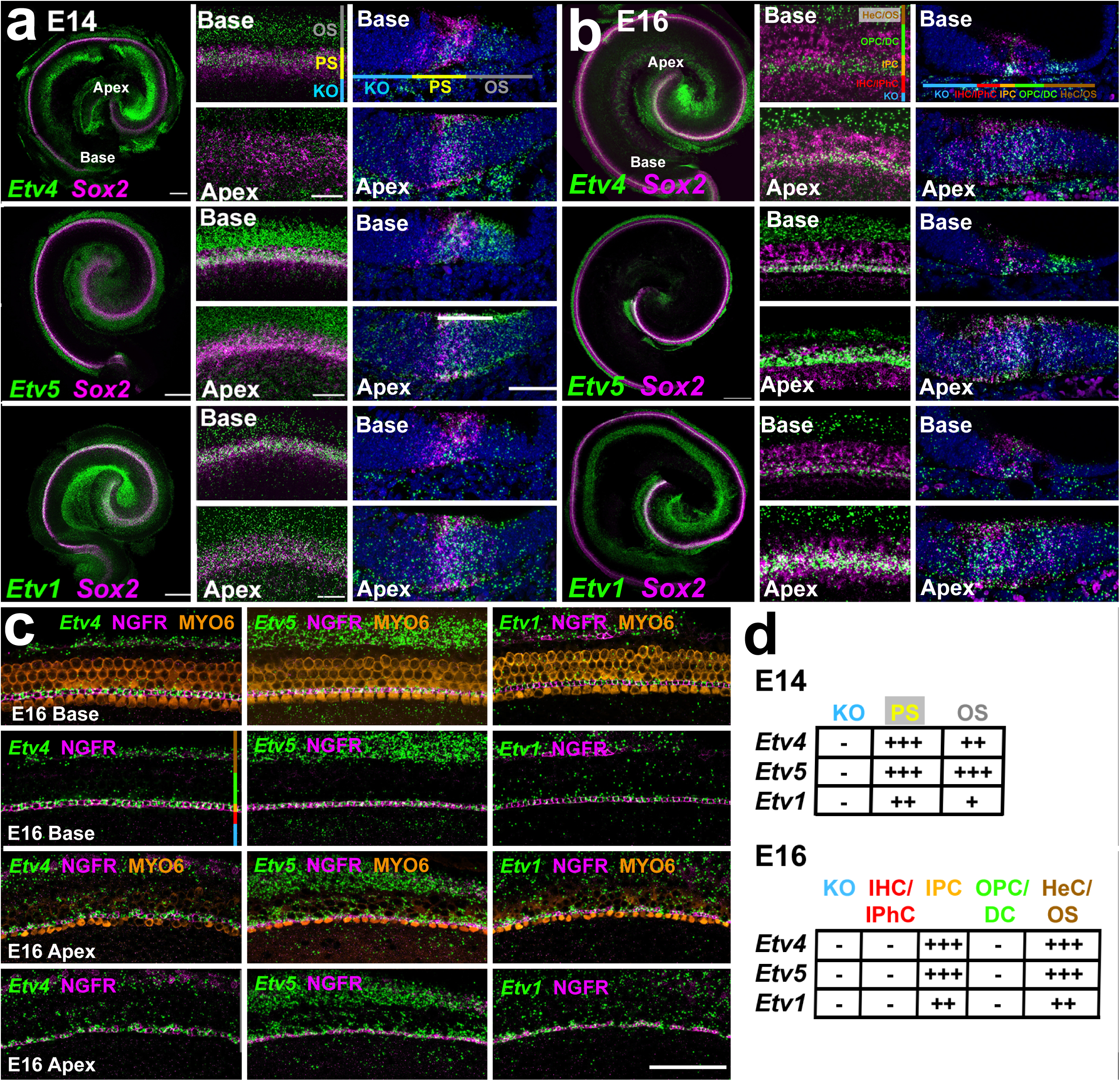
Expression of *Etv* genes during cochlear development. **a.** smFISH for *Sox2* (as a prosensory (PS) marker) and the indicated *Etv* genes in whole mounts (left), surface views (middle) and cryo-sections of the developing cochlear duct at E14. The locations of different domains within the cochlear duct are illustrated in the upper panels (refer to Fig. 1a). At E14, all three *Etv* genes are expressed along the length of the cochlea in a band that includes the *Sox2^+^*-domain and additional cells located in the outer sulcus (OS). In the more mature basal region, the band of *Etv* gene expression begins to split into two, one located within the PS domain and a second located in the OS. In the less mature apical region, all three *Etvs* are expressed in a single broader band that includes both the PS and OS. **b.** At E16, *Etv* gene expression persists along the length of the cochlea, but the pattern has become more refined. Expression in the PS domain has narrowed and in the basal region expression appears to be limited to one or two cell diameters. The position of *Etv* expression is consistent with the location of the developing IPCs. Laterally, broad expression of all three *Etvs* persists in the developing HeCs, located in the OS. **c.** Surface views of co-localization of *Etv4/5/1* with markers for HCs (MYO6) and IPCs (NGFR) at E16. In the more mature basal region of the OC, all three *Etv* genes precisely co-localize with NGFR. In the less mature apical region, expression of all three *Etv* genes is more diffuse but all three still show strong co-localization in the developing IPCs. **d.** Summary table for expression of each *Etv* gene in the regions indicated in panels a and b. Strong, diffuse expression in the PS becomes refined to just the IPCs while expression in the lateral OS/HeC domain is largely consistent across development. Abbr: KO, Kolliker’s organ. Scale bars in a, low magnification, 200 μm, high magnification surface views, 20 μm, Cryosections, 50 μm. Scale bar in c, 50 μm.,

At E16, whole mount images indicate continued expression of all three *Etv* genes along the length of the cochlea (Fig. 4b). Surface views show two bands of expression for each *Etv* gene, a narrow band located on the more medial side of the *Sox2*^+^-prosensory domain and a wider band in the region of developing HeCs (Fig. 4b, middle column). Histological sections confirm the refinement of *Etv* expression to a narrow band of cells within the developing OC and a wider band in the HeC/OS region (Fig. 4b, right column). Within the *Sox2*^+^-domain, expression of each *Etv* gene appears to correlate with the location of the developing IPCs.

At E18, while expression of all three *Etv* genes persists along the length of the cochlear spiral, expression within the *Sox2*^+^-prosensory domain has narrowed to one or two cell diameters (Suppl. Fig. 7a). A second band of expression in the HeC/OS region persists as well. To determine whether *Etv* gene expression within the developing OC is restricted to developing IPCs, smFISH for each *Etv* gene was combined with immunolabeling for markers for HCs (MYO6) and IPCs (NGFR) in E16 and E18 cochlear whole mounts (Fig. 4c and Suppl, Fig. 7b). Results indicate co-localization of each *Etv* gene and NGFR in the basal region of the cochlea at both E16 and E18. Similar co-localization is present in the apical region of the cochlea at E18. However, in the less mature apical region at E16 while *Etv* gene expression still co-localizes with NGFR, diffuse expression is also present in the region of developing OHCs, OPCs and DCs. A summary of *Etv* gene expression (Fig. 4d) indicates initial broad expression of all three *Etv* genes in both the prosensory domain and the lateral OS domain. As development continues, expression in the prosensory domain gradually refines to just the IPCs. *Etv* expression also persists in the HeC/OS region.

### Deletion of *Etv4/5/1* disrupts IPC development and OHC patterning

The results described above suggested that *Etv4*, *Etv5*, and *Etv1* are expressed in developing IPCs, that Etv4/5 regulons are probably active in developing IPCs, and that deletion of these genes is likely to perturb development of IPCs. To determine the specific roles of Etv signaling in IPC development, we sought to eliminate Etv signaling in the developing cochlea. Because of the overlap in expression of *Etv4*, *Etv5* and *Etv1* within the cochlea and the reported functional redundancy between *Etv* genes in other systems^44–47^, we generated mice with various tissue-specific deletions of *Etv4*, *Etv5* and *Etv1,* including animals lacking all three *Etv* genes. To achieve these phenotypes, a germline deletion line of *Etv4*, which is viable^48^, was combined with conditional alleles for *Etv5* and *Etv1*^47,49^. Deletion of the E*tv5^flox^* and *Etv1^flox^*alleles was driven by an *Emx2^cre^* allele which has been shown to be broadly expressed in the developing OC by E12^50,51^. Animals with multiple combinations of *Etv* deletions were collected at E18 because of likely neonatal lethality.

HCs and IPCs were labeled using anti-MYO6 and anti-NGFR respectively^27^. In control cochleae a single row of IHCs and three rows of OHCs were separated by a single row of IPCs in basal, middle, and apical turns (Fig. 5a, left column). Cochleae from animals lacking up to five of the six *Etv* alleles showed largely normal development except for occasional patterning defects (Fig. 5a, middle column and Suppl. Fig. 8). In contrast, *Etv* Triple Knock-Out (KO) cochleae showed significant defects in development. In particular, NGFR expression was almost completely absent throughout the length of the OC (Fig. 5a, right column). In the basal turn, additional, possible ectopic HCs were observed (Fig. 5a, right column, arrows) while in the apical region of the OC, OHC development appeared delayed and overall cellular pattern was disrupted. To determine whether the absence of NGFR expression in IPCs represented a specific effect on expression of this gene or a more global effect on IPC development, expression of a second IPC marker, NPY, was also examined. In controls, NPY was expressed in all IPCs (Fig. 5b, left panel). Consistent with the results for NGFR, NPY labeling was almost completely absent from the IPC region in *Etv* Triple KOs (Fig. 5b, right panel). In contrast with the results for NGFR, IPC expression of NPY was also perturbed in *Etv* Double KOs (Fig. 5b, middle panel). In addition to the lack of expression of NGFR and NPY in IPCs, in some images from the basal region of the cochlea, additional OHCs appeared to be present sometimes forming a partial fourth row (numbered in Fig. 5b right column). To examine this possibility more specifically, the density of OHCs and IHCs was determined for control and *Etv* Triple KOs at different positions along the OC. Despite the presence of ectopic OHCs in the basal OC region, OHC density was unchanged in the basal region of *Etv* Triple KO cochleae. However a significant decrease in OHC density was present in the mid and apical OC regions (Fig. 5c and Data File 4). To determine whether changes in cochlear length could contribute to the changes in OHC density, overall cochlear lengths were determined. No change in length was observed between control and *Etv* Triple KOs (not shown). (not shown). Subtle changes in OHC patterning were also observed in the basal region of *Etv* Triple KOs (Fig. 5a). To characterize these changes the spatial relationships of OHCs in the basal region of the OC were quantified (see Methods). In controls the spatial relationships of second row OHCs were highly ordered with cells forming a regular lattice (Fig. 5d, radar plot, see Methods for details). In contrast, in *Etv* Triple KOs, OHC spatial relationships were perturbed leading to significant changes in the crystalline organization of OHCs.

**Figure 5.**
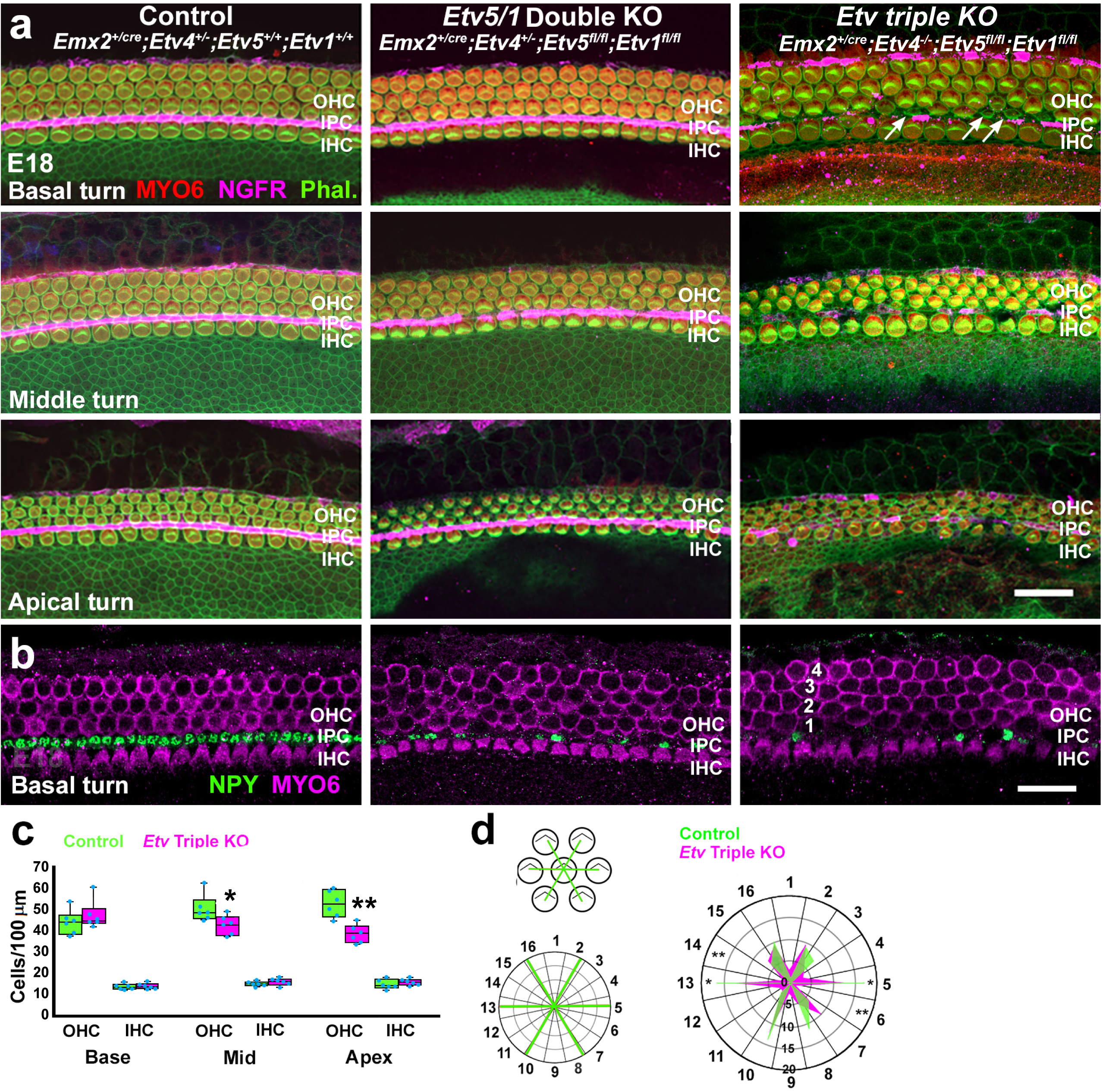
Deletion of *Etv4/5/1* leads to defects in IPC and OHC formation and OHC patterning. **a.** Surface views of the indicated regions of the OC in control, *Etv5/1* double KO and *Etv* Triple KOs at E18. In control a single row of IHCs is separated from three rows of OHCs by an ordered row of IPCs (NGFR in magenta). *Etv* double KO cochleae appear normal although some subtle patterning defects may be present in the apical OHC region. In *Etv* Triple KO cochleae, IPC development appears disrupted, NGFR expression is absent in most IPCs, and cellular development and patterning appears significantly disrupted in the apical region. In addition, small ectopic hair cells may be present in the basal turn in Etv triple Kos (arrows). **b.** Expression of a second IPC marker, NPY (green), is downregulated in both *Etv* double and Triple KOs. **c.** Quantification of HC density at different positions along the OC in control and *Etv* Triple KOs. Boxes indicate 1^st^ and 3^rd^ quartiles, interior line in each box indicates the mean, blue dots indicate individual data points. A significant decrease in OHC density is present in the middle and apical regions of the cochlea in *Etv* Triple KOs. **d.** Characterization of OHC patterning. The position of the six nearest OHCs were determined for OHCs located in the second row. Those positions were then plotted into 16 radial bins. The radar plot compares the frequency of OHCs in each radial bin in the basal region of the OC from control and *Etv* Triple KOs. Overall, *Etv* Triple KOs showed greater variability in the patterning of OHCs. Asterisks indicate sectors with significant changes between control (green) and *Etv* Triple KOs (magenta). * p<0.05; **, p<0.01. Scale bar in a (same in b) 20 μm. Abbr. Phal.: Phalloidin.

### Deletion of *Etv4/5/1* alters development of OPCs, and OHCs

In addition to defects in IPC development, our CellOracle analysis also predicted enhanced development of OPCs/DCs and HCs in the absence of Etv signaling. To examine these potential changes, cochleae from control and *Etv* Triple KO animals were labeled with anti-PROX1, a marker of all lateral supporting cell nuclei^52^ and anti-CD44, which, within the OC, is specific for OPCs^53^. It should be noted that CD44 also labels HeCs, but HeCs do not express PROX1. Therefore, PROX1^+^/CD44^+^ cells can be classified as OPCs. In control cochleae at E18, PROX1^+^-nuclei in the basal region form five ordered rows (Fig. 6a, left column). The medial-most row of PROX1^+^-nuclei (bottom of the panel) represent the IPCs which have developed a distinctive oblong nuclear shape. The row of PROX1^+^-nuclei just adjacent to these are the OPCs which also express CD44 in the basal region of the OC (Fig. 6a,a’, left panel). The lateral-most three rows of PROX1^+^-nuclei are the three well-ordered rows of DCs. In the less mature apex of control cochleae, IPC PROX1^+^-nuclei have not yet developed the oblong shape observed in the base and the more lateral rows of PROX1^+^-nuclei are not as highly organized (Fig. 6a).

**Figure 6.**
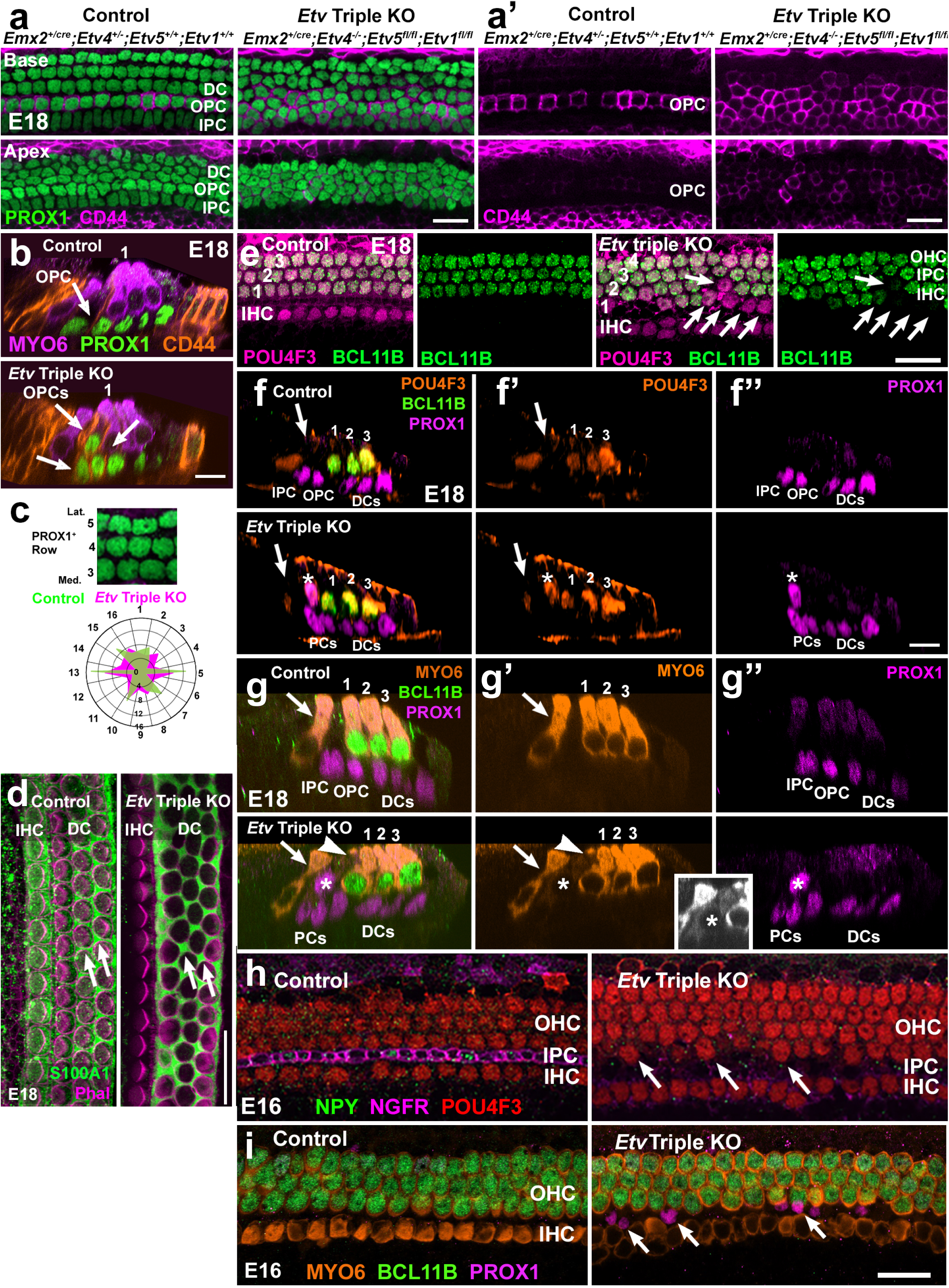
Deletion of *Etv4/5/1* alters OPC, DC and HC development and patterning. **a.** At E18, PROX1^+^-IPCs, OPCs and DCs are arranged in ordered rows in the basal region of the control cochlea. In the less mature apical region (lower panel) the lateral rows of DCs are not completely aligned yet. In contrast, in an *Etv* Triple KO cochlea overall cellular alignment is disrupted in both the basal and apical regions. **a’.** In a control cochlea, OPCs (CD44^+^, magenta) form a single ordered row in the basal region but are only weakly CD44^+^ in the apex. In an *Etv* Triple KO cochlea multiple rows of CD44^+^-OPCs are present in the basal and apical regions. **b.** Cross-sectional Z-stacks of the OC from the basal region of a control and an *Etv* Triple KO. In the control, the single CD44^+^/PROX1^+^-OPC (arrow) is located adjacent to the first OHC (numbered). In the *Etv* Triple KO, three OPCs (arrows), are intermixed with the first two rows of OHCs (numbered). **c.** The positions of the six nearest PROX1^+^-nuclei were mapped relative to fourth row PROX1^+^-cells (see text for details). The radar plot illustrates results for control and *Etv* Triple KOs. In control OCs, the spatial alignment of PROX1^+^-DCs is regular (green profile). In contrast, in *Etv* Triple KOs spatial patterning is disrupted (magenta). **d**. Surface view of the OC in control and Etv Triple KOs at E18 labeled with anti-S100A1, a marker of IHCs, PCs, and DCs, and phalloidin. In controls, DCs (arrows) form dumbbell shapes between OHCs. Although there are some patterning defects in the *Etv* Triple KO OC (arrows), overall expression of S100A1 appears comparable to control. S100A1-labeling of IHCs was also present in Etv Triple KOs (not shown). **e.** Surface views of an E18 control OC illustrating all HCs (magenta) and OHCs (POU4F3^+^ (magenta) and BCL11B^+^ (green)) and a similar view from an *Etv* Triple KO showing an over-production of HCs in the lateral region of the OC(numbered), many of which are negative for BCL11B (arrows). **f-f’’.** Cross-sectional Z-stack views of the OC from a control and an *Etv* Triple KO at E18. In the control, a single IHC (arrow), three OHCs (numbered), a single IPC and OPC, and three DCs are present. In an *Etv* Triple KO, an ectopic PROX1^+^-HC (asterisk) is present in the pillar cell region. **g-g’’.** Cross-sectional Z-stack views as in g but labeled with MYO6 instead of POU4F3. In the *Etv* Triple KO, a PROX1^+^/MYO6^+^-cell is present in the PC region of the OC (arrowhead and asterisk). Inset: Image of the ectopic hair cell (asterisk) in gray scale with increased contrast for improved detection of MYO6 labeling. **h.** Surface views of the basal region of the OC from animals with the indicated genotypes at E16. In controls a single row of NGFR^+^-IPCs (magenta) is already present. Some IPCs are also positive for NPY (green). In contrast, in an *Etv4/5/1* Triple KO only minimal expression of NGFR is present while NPY expression is absent. In addition, ectopic HCs (arrows) are present in the pillar cell space. **i.** Surface view of the basal region of the OC from control and an *Etv* Triple KO at E16. In the control a single row of MYO6^+^-IHCs and three rows of MYO6^+^/BCL11B^+^-OHCs are present. In an *Etv* Triple KO only three rows of OHCs are present by comparison with the four rows present in some regions at E18 (Fig. 5). Some PROX1^+^-nuclei are present in the HC nuclear layer (arrows) but are negative for HC markers. Scale bar in a,b,e,f,g (same in h), and j (same in i), 20 μm.

Visualization of CD44 expression indicates only faint expression in the row of developing OPCs (Fig. 6a’, left column). In contrast with controls, in the basal region of *Etv* Triple KOs, the medial most row of PROX1^+^-nuclei have not developed an oblong phenotype and the overall level of cellular organization is clearly disrupted (Fig. 6a, right panel). In addition, expression of CD44 is not confined to a single row of cells and instead forms approximately 3 poorly organized rows (Fig. 6a,a’, right column), suggesting an increase in the number of OPCs.

Consistent with this possibility, multiple rows of PROX1^+^/CD44^+^-cells are present in the apical region of the OC in *Etv* Triple KOs as well. The increased number of cells and overall level of CD44 expression in the *Etv* Triple KOs suggest premature or enhanced development of OPCs, a result that is consistent with the CellOracle analysis (Fig. 3a). To confirm the presence of ectopic OPCs in *Etv* Triple KOs, Z-stack images from the basal regions of control and *Etv* Triple KOs were examined. In contrast with controls which contain only a single PROX1^+^/CD44^+^ OPC (Fig. 6b, arrow), results from *Etv* Triple KOs indicate multiple PROX1^+^/CD44^+^-cells in the region between the IHC and first row OHC (Fig. 6b, lower panel, arrows). Whether these are cells that converted from an IPC or DC phenotype to become OPCs or are cells that have a mixed phenotype cannot be determined at this time. The images of PROX1-labeled cells in *Etv* Triple KOs suggested possible changes in the number of PROX1^+^-cells and a disruption in overall cellular patterning. To quantify these changes the density of PROX1^+^-cells and their cellular patterning was assessed as described for OHCs in the previous section. Cell counts indicated no significant changes in the density of PROX1^+^-cells in any region of the OC (Data File 4). For the cell arrangement assay, PROX1^+^-cells located in the fourth row along the medial-to-lateral axis were utilized. Results indicate a marked change in the crystalline alignment of PROX1^+^-cells in *Etv* Triple mutants (Fig. 6c). To determine if the absence of *Etv4/5/1* altered DC development, E18 cochleae were labeled with anti-S100A1 which labels IHCs, PCs and DCs (Fig. 6d). Results indicated comparable expression of S100A1 in DCs in *Etv* Triple KOs although some indications of patterning defects were evident (Fig. 6d, arrows).

While quantification of HC density in the basal regions of *Etv* Triple KO cochleae did not indicate a significant increase, ectopic rows of HCs were observed in some samples (Fig. 5b). To determine whether these cells represented ectopic OHCs, as have been reported in other mutants with defects in IPC development^54,55^, cochleae from control and *Etv* Triple KOs were labeled with anti-POU4F3, a HC marker, and anti-BCL11B, a specific marker of OHCs^7,56^. In contrast with controls in which all HCs in the lateral region were positive for BCL11b, some HCs in the lateral OC region in *Etv* Triple KO cochleae were BCL11B negative (Fig. 6e, arrows). To determine whether these ectopic HCs were developing from PROX1^+^-cells, expression of PROX1, POU4F3 and BCL11B was localized in Z-stacks of control and *Etv* Triple KOs. Expression of PROX1 in a developing HC would suggest an ectopic HC forming due to the absence of Etv signaling. In controls, a single IHC, three OHCs, two pillar cells and three DCs can clearly be visualized (Fig. 6f-f’’, upper row). All PROX1^+^-nuclei are located along the basement membrane of the OC. In contrast, in *Etv* Triple KOs a fourth HC (Pou4f3^+^, asterisk) is present in the pillar cell region (Fig. 6f-f’’, bottom row). The nucleus for this HC cell is positive for PROX1 but is located at the level of other HC nuclei (Fig. 6f’’) suggesting that it has developed from a PROX1^+^-supporting cell. To confirm the identity of these misplaced PROX1^+^-cells, additional samples were labeled with an alternative HC marker, MYO6, which labels the HC cytoplasm. While weakly expressed, MYO6-labeling was also present in ectopic PROX1^+^-HCs (Fig. 6g). In all cases, ectopic HCs were negative for expression of BCL11B, suggesting they are either immature or, possibly, IHCs.

The results described above indicated significant changes in determination of cell fates in the OC in the absence of *Etv4/5/1* signaling. To determine the timing of the onset of those changes, cochleae from control and *Etv* Triple KOs were collected at E16 and labeled with markers for IPCs, HCs and OHCs. In controls, developing IPCs are already NGFR^+^ but show limited expression of NPY (Fig. 6h, left panel). In *Etv* Triple KOs, NGFR expression is nearly undetectable and no NPY expression is observed (Fig. 6h, right panel). Moreover, some ectopic HCs are present in the pillar cell region (arrows in 6h, right panel). Consistent with the observation of ectopic HCs, localization of PROX1 at E16 indicates the presence of misplaced PROX1^+^-cells in the pillar cell space (Fig. 6i). These results suggest that, in the absence of Etv signaling, OC development is perturbed prior to E16.

### *Etv* expression is down-regulated in *Fgfr3^-/-^*mutants

Based on the similarity in the cochlear phenotypes between *Etv*4/5/1 Triple KOs and *Fgfr3^-/-^*mutants^31,54,57^ and previously demonstrated interactions between Etvs and Fgfs in other systems^45,47,58,59^, we next examined changes in expression of *Etvs* in cochleae from *Fgfr3^-/-^*mutants at E14, E16 or E18. Results indicate similar patterns of expression for *Etv4, Etv5* and *Etv1* in control and *Fgfr3^-/-^* mutant cochleae at E14 (data not shown). However, at E16 and E18, expression of all three *Etv* genes is down-regulated in IPCs in *Fgfr3^-/-^* mutants (Fig. 7a and Suppl. Fig. 9). Down-regulation was more pronounced in the basal region of the cochlea, possibly reflecting a temporal progression in *Etv* expression along the cochlear developmental axis. Expression of *Etv* transcripts in the lateral HeCs was unchanged in *Fgfr3^-/-^* mutants. These results suggest that while initial expression of *Etv 4/5/1* does not require activation of Fgfr3, maintenance of *Etv* expression in the developing IPCs is dependent on Fgfr3 signaling. In contrast, expression of *Etvs* in HeCs seems to be independent of Fgfr3 signaling.

**Figure 7.**
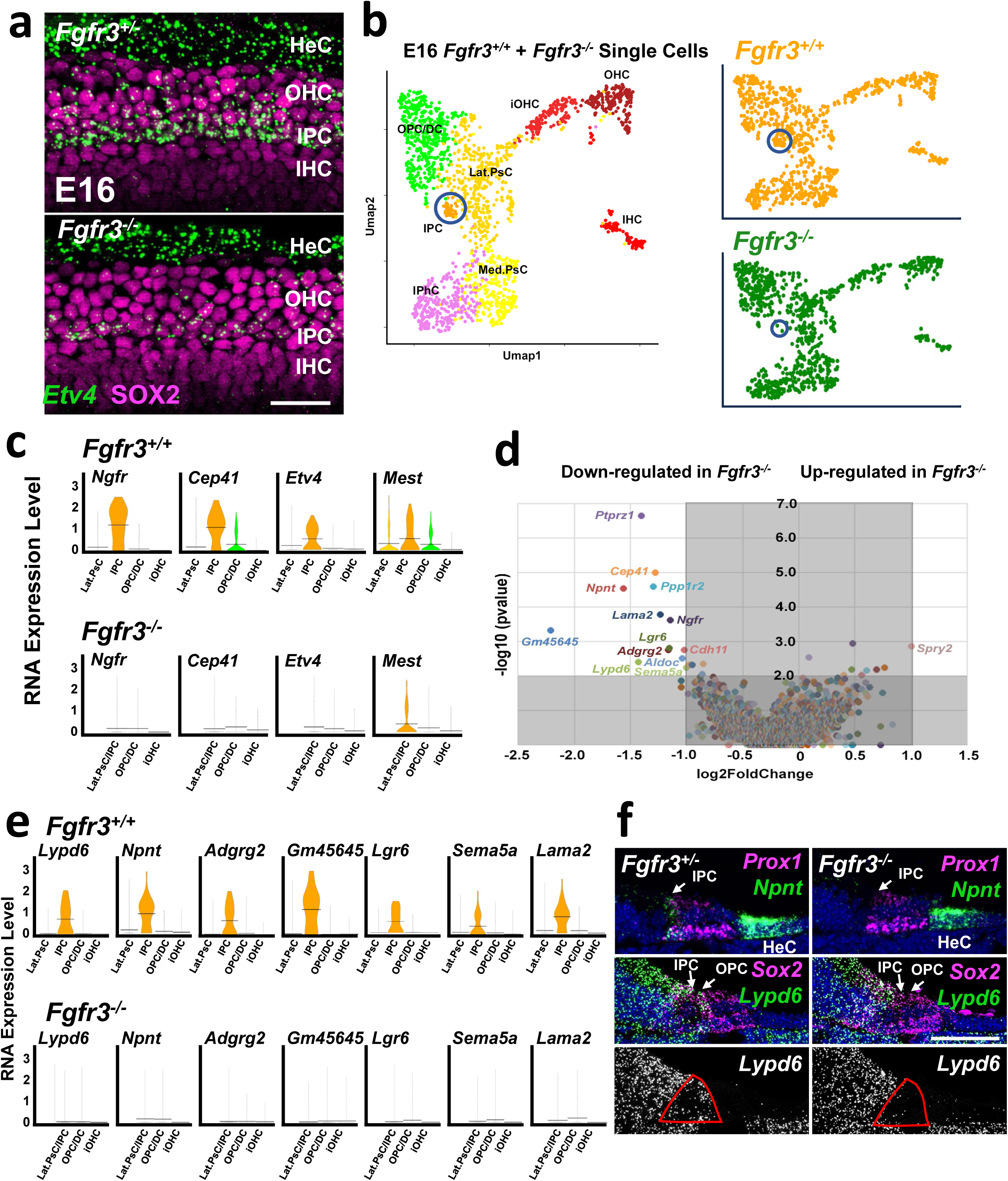
Analysis of cochleae from *Fgfr3^-/-^*mice at E16. **a.** Surface views of smFISH analysis of expression of *Etv4* (green) and SOX2 (magenta) in the basal region of the cochlea from *Fgfr3^+/-^* and *Fgfr3^-/-^* mice. In the *Fgfr3^+/-^* cochlea the single row of IPCs is positive for *Etv4.* A second region of *Etv4* expression is present in the HeCs. In the *Fgfr3^-/-^* sample, while *Etv4* expression is still present in the HeCs, expression in the IPC region is absent. **b.** Left: combined scRNAseq UMAP for *Fgfr3^+/+^*and *Fgfr3^-/-^* cells. Cluster identities are marked including IPCs (circled). Right: the same plot as on the left but separated based on genotype. There is consistent overlap for all cell types except IPCs (circled). **c.** Violin plots for expression of four predicted Etv target genes (Fig. 3) in lateral OC cell types from *Fgfr3^+/+^*and *Fgfr3^-/-^*single cells. Because only two IPCs were recovered from *Fgfr3^-/-^* cochleae, these cells were combined with Lateral Prosensory Cells. *Ngfr*, *Cep41* and *Etv4* are not expressed in *Fgfr3^-/-^*lateral cell types while *Mest* shows decreased expression. **d.** Volcano plot illustrating results of pseudobulk analysis of changes in gene expression of lateral cells between *Fgfr3^+/+^* and *Fgfr3^-/-^*(see Methods for details). Thirteen genes show a significant decrease, and one gene shows a significant increase. **e.** Violin plots for seven of the thirteen significantly decreased genes from d. All are specifically expressed in IPCs, and all are absent in cells from *Fgfr3^-/-^* mice. **f.** Cochlear cryosections at E16 illustrating expression of *Lypd6* and *Npnt* in *Fgfr3^+/-^*and *Fgfr3^-/-^*using smFISH. The prosensory domain is marked by expression of *Prox1* for *Npnt* or *Sox2* for *Lypd6* in magenta. In the control, *Npnt* is expressed in IPCs and Hensen’s cells (HeC) but in the *Fgfr3^-/-^* cochleae expression in the IPC is absent. *Lypd6* is expressed in both inner and OPCs but is also expressed in more medial regions of the epithelium. Lower panels show just *Lypd1* expression in gray scale. The red outline indicates the position of the IPCs and OPCs. Scale bar in A, 50 μm, Scale bar in F, 50 μm.

### Changes in gene expression in *Fgfr3^-/-^* IPCs

Based on the results described in the previous sections, we wanted to identify potential early downstream targets of Fgfr3/Etv signaling in IPCs. To do this, scRNAseq was performed on cochlear cells collected from E16 control (*Fgfr3^+/+^*) and *Fgfr3^-/-^* mutant cochleae. *Fgfr3^-/-^*mutants were selected because of the challenges involved in generating *Etv* Triple KOs. Following collection and initial analysis, PsC/OC cells were isolated bioinformatically and then merged into a single data set. We identified eight clusters; IHC, iOHC, OHC, iMed.PsC, Med.PsC, Lat.PsC, OPC/DC and IPC, based on similarity with the previously described dataset (Fig. 7b and Data File 2). When the data sets were separated based on genotype, cells mapping to each cluster were present in both genotypes, however the number of IPCs originating from the *Fgfr3^-/-^*collections was reduced to two (Fig. 7b, circled). To determine whether Etv downstream signaling is altered in the absence of *Fgfr3*, the expression of four putative Etv target genes that were completely or nearly IPC-specific (Fig. 7c) was examined. Because so few IPCs were captured from *Fgfr3^-/-^* mutants, Lat.PsCs, the precursors for the IPCs, were combined with the IPCs. Results indicate either a complete, or near complete, loss of expression for all four putative Etv targets including *Etv4* (Fig. 7c). Putative Etv target genes that were not IPC-specific showed variable levels of change (Suppl. Fig. 10). Similarly, OPC/DC and HC specific genes showed minimal changes in expression (Suppl. Fig. 11) suggesting that the initial effects of Fgfr3/Etv signaling are restricted to developing IPCs.

To identify other potential regulators of Fgfr3/Etv signaling, we performed a pseudobulk RNAseq analysis^60^ to compare overall changes in gene expression between control and *Fgfr3^-/-^*cochlear cells (Fig. 7d and Data File 5). Using a fold change cutoff of 2.0 (log2 = 1.0) and a - log10 p value of 2.0 (p < 0.01) thirteen genes were found to be significantly down regulated. Of these, nine genes were already known to be or were determined to be IPC specific (Fig 7e), and all showed a complete absence of expression in the Lat.Psc/IPC cluster from *Fgfr3^-/-^* cells. Only one gene, the Fgf antagonist *Spry2*, showed a marginally significant upregulation in *Fgfr3^-/-^* cells. To confirm the results of the single cell analysis, expression of two genes that were significantly down-regulated in the pseudobulk analysis, *Npnt* and *Lypd6* were examined by smFISH in control and *Fgfr3^-/-^* cochleae at E18. Results indicated an almost complete absence of *Npnt* and *Lypd6* expression in *Fgfr3^-/-^* mutant cochleae (Fig. 7f). Interestingly, smFISH also indicated limited expression of *Lypd6* in OPCs. While *Lypd6* expression was not detected in the scRNAseq dataset, the smFISH results suggest that this could be a new marker for OPCs. Upregulation of *Spry2* was not confirmed. These results demonstrate a crucial role for Etv signaling in IPC development as well as identifying novel down-stream regulators of pillar cell identity.

## Discussion

The OC is a highly derived cellular structure found only in placental mammals^6,61^. The overall organization includes a number of specialized cell types and components that have evolved to mediate increased frequency range and discrimination. Among these are the IPCs and OPCs that combine to form the tunnel of Corti, a fluid filled space extending along the basal-to-apical axis of the cochlea that is thought to play a role in cochlear stability and intracellular transport^62^. In the adult OC, IPCs contain elongated microtubules that confer stiffness and stability to these cells^63^. But during development, IPCs express a number of genes, such as *Ngfr* and *Npy*, that have no known link to cellular architecture. Rather, expression of these genes suggests potential additional roles for IPCs in some aspects of neuronal signaling, patterning or survival. The results presented here identified several novel genes that are exclusively expressed in IPCs.

These include *Npnt, Cep41, Crispld1, Lama2, Lypd6, Adgrg2* and one predicted gene, *Gm45645*. The known roles of these molecules are fairly disparate, with NPNT acting as a receptor for INTEGRIN α8β1^64,65^, CEP41 stabilizing microtubules^66,67^, LAMA2 playing a role in regulation of basement membrane formation and muscular dystrophy^68,69^ and LYPD6 acting as a regulator of cholinergic signaling^70,71^. Some of these processes, such as regulation of basement membrane formation, interactions with the basement membrane and regulation of microtubules are all consistent with known aspects of pillar cell development and/or function, but others, including synaptic signaling, are more difficult to interpret. Determination of the specific roles of these genes in pillar cells will probably require focused analyses of deletion phenotypes.

Despite an incomplete understanding of the intracellular factors that mediate IPC development, our understanding of the developmental interactions that specify IPCs is becoming clearer ^31,54,55,57,72–75^. Following terminal mitosis, cochlear PsCs rapidly become sub-divided into medial and lateral PsCs. At this stage, *Etv4, 5* and *1* are broadly expressed both within the PsC domain and in cells located in the more lateral OS region. The factors that regulate the initial expression of *Etv4/5/1* in the cochlear duct have not been determined but recent results have shown decreased expression of all three Etvs in cochleae from *Fgfr20^-/-^* mice^38^. As development continues, *Fgfr3* is upregulated in the Lat.PsCs and developing IHCs located adjacent to the medial edge of the Lat.PsC domain upregulate expression of *Fgf8*. Expression of *Fgfr3* is indicative of the potential for many, if not all, cells within the Lat.PsC population to develop as IPCs, however, activation of FGFR3 is required for IPC development. Under normal circumstances, only the row of cells located directly adjacent to the IHCs receive a sufficiently high dose of FGF8 to develop as IPCs. If *Fgf8* or *Fgfr3* are deleted or if activation of FGFR3 is inhibited, IPCs fail to form. In contrast, if the Fgf pathway antagonist SPRY2, which is expressed in much of the Lat.PsC domain, is removed or the concentration of FGF is increased, cells that would have developed as OPCs or DCs will be converted to IPCs. The results presented here demonstrate that a key step in that process is FGFR3-dependent maintenance of *Etv4/5/1* expression in developing IPCs. As discussed above, continued expression of *Etv4/5/1* upregulates a number of initial IPC genes which play roles in cellular structure, interactions with the basement membrane and, possibly, axon guidance. While not tested in this study, it seems likely that forced prolonged expression of ETV4/5/1 in additional Lat.PsCs would lead to the formation of additional IPCs. A final important factor in IPC development is activation of the transcriptional activity of β-Catenin (β-Cat). Targeted disruption of *β-Cat* using *Emx2^cre^* or even an IPC-specific *Npy^cre^*, induces a phenocopy of many of the aspects of the *Etv* Triple KO, in particular, an absence of IPCs and an overproduction of OPCs^76,77^. The specific interactions between the Fgfr/Etv and β-Cat signaling pathways are unclear. scRNAseq data from *β-Cat* mutant cochleae indicated a decrease in *Etv4* but no changes in expression of *Etv5, Etv1* or *Fgfr3*^76^, suggesting that β-Cat might function downstream of Fgfr3 or in a parallel pathway to specify IPC identity.

In addition to disruption in IPC development, deletion of *Etv4/5/1* has other significant effects on OC development. In particular, distribution and patterning of OHCs and PROX1^+^-supporting cells was altered and ectopic HCs were observed in the pillar cell region in the cochlear basal turn. These changes resulted in significant decreases in the density of OHCs in the mid and apical regions of the OC and downward, but not significant, trends in PROX1^+^-cellular density in the same regions. In contrast, the densities of both OHC and PROX1^+^-cells in the basal region of *Etv* triple mutants showed an upward, but not significant, trend. These results, along with the changes in the normally organized cellular patterning of the OC, are consistent with a potential perturbation in cellular migration and/or outgrowth during cochlear extension. Previous studies have demonstrated that defects in cellular migration result in a similar phenotype and changes in IPC shape and patterning were demonstrated in response to perturbation of non-muscle Myosin II. However, it is currently not possible to determine if all of the changes observed in *Etv* triple mutants are a result of the defect in IPC formation. All three *Etv* genes are initially expressed broadly in the developing lateral region of the OC suggesting a possible early role in the development of other OC cell types. Additional experiments using an IPC-specific cre line, such as *Npy^cre^* would be required to begin to separate the roles of *Etv* genes in IPC versus other OC cell types.

One aspect of the phenotype observed in Etv triple KOs, the formation of ectopic HCs in the basal region, is consistent with changes in cell fate that were reported for cochleae with disruptions in either Fgfr3 or Hey2 signaling^54,55,57^. These findings, along with the results presented here, are consistent with the hypothesis that if the inductive signal for IPC development is removed or inhibited, some Lat.PsCs will develop as ectopic HCs.

Some insights regarding potential Fgfr3-independent roles of Etvs can be gained by examining the phenotypes in other pillar cell mouse mutants. While the similar IPC phenotype^54,57^ and the demonstration that expression of *Etv4/5/1* is dependent on *Fgfr3* clearly links the roles of *Fgfr3* and *Etv4/5/1* in OC development, there are some notable differences between the two mutant phenotypes. In particular, OPCs have been reported to be absent in *Fgfr3^-/-^* mutants and the number of HCs in the middle and apical region of *Fgfr3^-/-^* cochleae is increased^53^ rather than decreased as is the case in *Etv4/5/1* Triple KOs. Further, the overall level of disruption in cellular patterning, in particular in the middle and apical regions of the OC, appears to be higher in the *Etv4/5/1* Triple KOs by comparison with *Fgfr3^-/-^* cochleae. These results suggest that the initial, Fgfr3-independent expression of *Etv4/5/1* throughout the developing OC may play a role in overall cellular patterning and, possibly, HC or DC fate specification.

Based on the results of lineage-tracing experiments for the development of the OC^11^, HeCs were excluded from the OC Sensory Dataset. However, smFISH indicated expression of all three *Etv* genes in the HeCs. *Etv* expression in the HeCs is not dependent on FGFR3 activation as *Etv* expression was still present in *Fgfr3^-/-^*cochleae. Moreover, while not examined specifically, deletion of *Etv4/5/1* did not appear to have a significant impact on HeC development. CD44, a marker of both OPCs and HeCs, appeared largely normal in *Etv4/5/1* Triple KO cochleae (Figure 6b, f). Determination of the role of Etvs in HeCs may require a more targeted deletion using a HeC-specific driver, as changes in development of the OC could have secondary effects on HeC formation.

In addition to the identification of *Etv4/5/1* as crucial regulators of IPC development, the OC Sensory Dataset presented in this study has the potential to provide insights regarding the development of other OC cell types. A recent study used scRNAseq data from E12.5 and E14.5 cochlear ducts to suggest roles for Shh and RA in development of tonotopy, however, that study grouped all the cochlear floor cells together^21^. Here, we focused on the same developmental time period, but by including cells from E16, we were able to generate developmental trajectories extending from the prosensory cells through several novel transitional cell types to most of the known cell types within the OC. Of particular interest may be the Lat.PsC population. Recent lineage tracing results have confirmed strong clonal relationships between cells located in the lateral compartment of the OC^11^, suggesting a specific developmental bias within the Lat.PsC. cell population. This population of cells is known to express a number of unique markers, such as PROX1, *Fgfr3*, *Tgfbr1*, *Fzd9*, CDH1, and possibly *Emx2*, beginning as early as E14^15,41,72,78,79^, but results to date have not demonstrated that deletions of any of those genes leads to a complete absence of lateral OC cell types. The SCENIC analysis presented here identified a number of novel candidates, such as Rxrγ, Foxg1, and Tbx21 which have not been previously examined for roles in lateral prosensory cell specification.

The development of novel analysis packages that incorporate the growing wealth of single cell RNAseq and ATAC data have the potential to rapidly identify candidate genes for the specification of specific cochlear cell types. In this study, we applied CellOracle^39,40^, which combines scRNAseq and single cell or bulk ATACseq data to build predictive regulatory networks for cellular specification and state changes, to our OC Sensory Dataset. CellOracle correctly predicted the primary effects of deletion of several transcription factors that are known to regulate development of HCs, or SCs. Similarly, the simulated deletion of *Etv4* predicted defects in development of IPCs and an enhancement in the development of OPCs that were supported by our analysis, although it should be noted that the phenotype was only obtained once *Etv5* and *Etv1* were also deleted. However, for each of the predicted gene deletions, the results of the CellOracle analysis also predicted enhancements of cell type development that were not confirmed. There are several caveats to consider. First, because CellOracle makes predictions based on changes in gene expression and/or chromatin accessibility, it may be the case that the predicted enhancements are occurring. A specific analysis of changes in gene expression or accessibility would be required to determine the extent of these possible changes. Moreover, because we used an existing CellOracle GRN data base that was built using ATAC data from adult tissues^80^ that did not include the inner ear, it is possible that some of the predictions may not be as accurate as they might be with the inclusion of embryonic cochlear ATAC data. Finally, it is important to note that the overall design of CellOracle cannot account for potential secondary signaling effects in which a change in the fate of one cell type might then alter the developmental trajectory of another cell type through undefined paracrine signaling. Overall, CellOracle seems well suited for predicting the primary, intracellular effects of transcription factor deletion but may not be as accurate for predicting secondary changes.

In conclusion, using a novel scRNAseq dataset for the early development of OC cell types, we identified the ETV4/5/1 transcription factors as likely regulators of IPC development. Generation of *Etv4/5/1* Triple KO mice confirmed the predicted effects of Etv genes in development of IPCs. In addition, analysis of cochleae from *Etv4/5/1* Triple KOs demonstrated additional roles for Etv4/5/1 in development of OHCs and lateral SC and in cellular patterning. Finally, scRNAseq analysis of *Fgfr3^-/-^* cochleae at E16 identified a group of genes that are downregulated in response to disruption of IPC development. These results should provide novel insights regarding the molecular pathways regulating the formation of IPCs and the tunnel of Corti.

## Methods

### Animals

All animals were housed in the Porter Neuroscience Research Center Shared Animal Facility, and were maintained following guidelines set by the NIH. The animal protocol was reviewed and approved by the NINDS/NIDCD Animal Care and Use Committee. Timed-pregnant CD1 females were obtained from Charles River Laboratories. *Etv1^flox^*, *Etv4^-^* and *Etv5^flox^* mice were obtained from MMRRC and maintained in a mixed genetic background^47–49^. *Emx2^Cre^* mice were obtained from Shinichi Aizawa at RIKEN Center for Developmental Biology and maintained in the C57BL/6J background^50^. *Fgfr3^+/-^* mice were maintained on C57BL6 background, as described previously^57^. Littermate controls were used for all comparisons with mutant phenotypes. Animals of both sexes were used.

### Tissue collection

Pregnant CD1 mice were euthanized at embryonic days 11, 12, 13, 14 or 16. Embryos were dissected out, staged for developmental time point, and their cochleae were isolated. Animals of both sexes were used. Cochleae were incubated for 10 to 15 min in 3 ml DMEM:F12 with 100 μl Thermolysin (5 mg/ml) and 5 μl DNase (1 mg/ml) at 37°C. Subsequently, the mesenchyme, roof of the cochlea, and lateral wall were carefully dissected away, and the epithelial floor of the cochlear duct was isolated. Cells were dissociated using pre-warmed papain (20 U/ml) for 30-40 minutes, with gentle trituration performed every 5 minutes. Following dissociation, papain was neutralized with an ovomucoid inhibitor. Cells were then passed through a 20 μm cell strainer, centrifuged at 300 g, and resuspended in PBS with 0.4mg/mL BSA and 0.2U/mL RNase inhibitor. Single dissociated cells were loaded onto the 10X Genomics Chromium Controller, where they were lysed, mRNA was reverse transcribed into cDNA, and sequencing libraries were prepared using Chromium Single Cell 3ʹ Reagents (v3.1) according to the manufacturer’s instructions. Libraries were sequenced using the Illumina NextSeq platform and mapped to the Ensembl GRCm38 mouse genome with Cell Ranger software (10X Genomics). A minimum of two collections were made at each developmental time point.

### Single cell RNAseq analysis

For the data set presented in this study, we combined the following newly obtained collections: four samples from E11, four samples from E12, three samples from E13, two samples from E14, and three samples from E16 with previously reported data sets obtained at E14 and E16^15^. The Cell Ranger output data was analyzed using Seurat v4^81^. For the analysis, cells with fewer than 500 unique genes or exceeding 20,000 RNA counts were removed. Cells with greater than 8% mitochondrial genes or more than 5% hemoglobin transcripts were also excluded from further steps. Each dataset was processed through normalization, scaling, and dimensionality reduction following the standard Seurat workflow. Seurat’s "ScaleData" function was used to regress out RNA counts and mitochondrial content scores. Principal components analysis (PCA) and uniform manifold approximation and projection (UMAP) were employed as non-linear dimensionality reduction techniques for visualization on a 2D UMAP plot, followed by unsupervised clustering analysis. Clusters of mesenchymal cells expressing *Twist1* or *Pou3f4*, as well as neuronal cells expressing *Nefl* or *Nefm*, were excluded from each dataset, retaining only cells from the cochlear duct. Datasets were integrated using Seurat RPCA, followed by the standard Seurat workflow for UMAP visualization and unsupervised clustering. Clusters were evaluated for quality and excluded if they showed expression of mesenchymal or neural genes. To obtain sensory epithelial progenitor cells and sensory epithelial cells, clusters of *Atoh1*-positive HCs and *Matn4*-positive inner phalangeal cells were isolated. Since *Tac1*-positive cells were predominantly located in the apical turn and formed a large cluster containing a substantial number of non-sensory epithelial cells, *Sox2*-positive *Tac1*-negative sensory epithelial clusters were specifically selected for further analysis. Datasets were reprocessed using the standard Seurat workflow for UMAP visualization and unsupervised clustering.

### RNA velocity analysis

Using the BAM files generated by the Cell Ranger alignment, we applied Velocyto (v0.17.16) to obtain loom files containing spliced and unspliced read count matrices^82^. Spliced and unspliced read count matrices for the cells included in the Seurat-generated dataset were extracted and integrated into the dataset using scVelo (v0.3.3)^83^. RNA velocity was inferred with UniTVelo (v0.2.5) without specifying root cells^33^. Using scVelo, we projected the estimated velocities onto the precomputed UMAP embeddings and displayed them as streams.

### Trajectory analysis

Trajectory analysis was performed using Slingshot and Monocle3. Slingshot (R package v2.10)^34^ was performed using UMAP as the dimensionally reduced input. The cluster containing the most immature prosensory cells, marked by the absence of *Cdkn1b* expression, was identified as the starting cluster. The end clusters (IPC, OPC/DC, IPhC, IHC, and OHC) were determined by their marker gene expression profiles. The TradeSeq (R package v1.16.0)^84^ analysis was conducted on Slingshot trajectories with a Kknots value of 8 to identify genes that change across pseudotime. To perform trajectory analysis using Monocle3(R package v1.3.1)^85^, the SeuratObject was converted into a CellDataSet (CDS). Trajectory analysis was performed by selecting a cell within the immature prosensory cell cluster as a root cell.

### Gene Regulatory Networks (GRN) analysis

GRN analysis was performed using pySCENIC^86^ and CellOracle^40^. To perform GRN analysis using pySCENIC and CellOracle, the SeuratObject was converted into a AnnData file. GRNBoost2, implemented in pySCENIC, was used to infer co-expression modules, identifying potential regulatory interactions between transcription factors and target genes. CisTarget was then used to prune indirect targets by leveraging the dataset generated by GRNBoost2 combined with cisTarget databases (mm10 500bpUp100Dw and TSS ± 10 kbp), thereby refining the co-expression modules into candidate regulons. AUCell was used to quantify the activity of each regulon within individual cells, generating regulon activity scores. The activity of a regulon within a specific cluster was evaluated using its regulon specificity score (RSS). In silico TF perturbation of gene regulatory networks (GRNs) was carried out using CellOracle (v0.18.0) and the results of the Monocle3 trajectory analysis. Using a pre-built GRN from mouse sc-ATAC-seq atlas data^80^, we generated an Oracle object and constructed cluster-specific GRNs for all clusters. Several network scores were then calculated. Active regulatory genes from the GRN were selected for further analysis. For each active regulatory gene, a simulated TF perturbation vector field was generated, and the perturbation score (PS) was calculated for each lineage.

### Determination of HC or PROX1^+^-cell density

Cochleae from E18 control and *Etv4/5/1* Triple KO animals were collected, labeled with a HC marker, typically anti-MYOSIN6 and/or anti-PROX1. Cochleae were then dissected and mounted for imaging. Images were obtained from the basal, mid and apical regions of the duct using a Zeiss LSM900 microscope. The length of the OC within each image and the number of HCs and/or PROX1^+^-cells was then determined using Zen Black. The number of HCs or PROX1+ cells was then divided by the length of the sensory epithelium in the image to generate cell density. This number was multiplied by 100 and then plotted. For quantification of hair cell density, 6 samples from 6 separate animals were analyzed for each genotype. For PROX1+ quantification, 3 samples from three separate animals were collected for each genotype. Significance was determined using two-way ANOVA and post-hoc Tuckey’s multiple comparison test.

### Determination of cellular patterning

Images of the lumenal surface of E18 cochleae from control and *Etv* Triple KOs were prepared as described above. For OHCs a six-pointed star was place over the center of the lumenal surface of each OHC in the second row and the six arms of the star were connected to the center of the lumenal surface of the six closest OHCs. Each star was than overlayed on a radial map divided into 16 equivalent sections encompassing 22.5° of arc per section and numbered 1 to 16. The frequency of individual arms in each section was determined for a minimum of 10 HCs per region per sample. Data were acquired for a minimum of 3 cochleae from 3 mice per genotype. For PROX1^+^-DCs, the same approach was used. PROX1^+^-DCs located in the second most lateral row were measured. Significant changes were determined using a T-test to compare changes in frequency of section occupancy.

### Single cell analysis of control and *Fgfr3^-/-^*cochleae

Four samples were dissected from *Fgfr3^−/−^* mice at E16 and three samples were collected from Fgfr3 wildtype (WT) mice at the same age (The WT samples were also included in the larger data set described above). Dissociation and single cell capture was as describe above. Next, Seurat was used to perform the analysis following the same procedure as described for CD1 mice above. Reciprocal principal components analysis (RPCA) was applied for integration. To investigate the differences in sensory epithelium between WT and KO, sensory cells were separated from the integrated dataset containing both WT and KO. UMAP visualization and clustering were then re-performed on this subset. Similar numbers of cells were obtained, on average from WT and KO cochleae, however fewer cells were identified as IPCs in the KO samples. All other supporting cell types were identified. To examine the differences in gene expression between WT and KO, the IPC and Lat.PsC clusters were combined. Pseudobulk analysis was then performed on these combined clusters using DESeq2 (version 1.47.3)^60^.

### Immunohistochemistry (whole-mount preparations)

Embryos were collected from timed pregnant females at specific embryonic timepoints, as indicated. Inner ears were dissected out from the skull and fixed in fresh 4% PFA for 1.5 h at room temperature. Fixed inner ears were washed in PBS overnight at 4°C. Cochlear roof, Reisner’s membrane and the tectorial membrane were subsequently dissected off to expose the sensory epithelium of the cochlea. Cochleae were blocked in PBS containing 10% normal donkey serum and 0.5% TritonX for 1 h at room temperature. Cochleae were incubated in primary antibody overnight at 4°C. Cochleae were then incubated in secondary antibodies in PBS for 1 h at room temperature. After secondary antibody incubation, the sensory epithelium was dissected and mounted on a glass slide. For NPY and BCL11B antibodies, antigen retrieval was performed prior to the initial blocking step as follows; samples were incubated in citrate buffer (10 mM citric acid, 0.05% Tween-20, pH 6) at 70°C for 20 minutes. The primary antibodies used were rabbit anti-MYOSIN6 (Proteus Biosciences 25-6791, RRID:AB_10013626 at 1:500 dilution), rabbit anti-NPY (Peninsula Laboratories T-4070, RRID: AB_518504 at 1:500 dilution), goat anti-NGFR (R&D Systems AF367, RRID:AB_2152638 at 1:500 dilution), goat anti-PROX1 (R&D Systems AF2727, RRID:AB_2170716 at 1:500 dilution), goat anti-SOX2 (R&D Systems AF2018, RRID:AB_355110 at 1:500 dilution), rat anti-CD44 (BD biosciences 550538, RRID: AB_393732 at 1:500 dilution), mouse anti-POU4F3 (Santa Cruz biotechnology, RRID: AB_2167543 at 1:200 dilution), goat anti-MYOSIN6 (Kelley lab custom made at 1:500 dilution), mouse anti-CTIP2 (BCL11B) (Abcam ab18465, RRID: AB_2064130 at 1:500 dilution), rabbit anti-NRP2 (Cell Signaling D39A5, AB_2155250 at 1:500), sheep anti-S100A1 (R&D Systems, AF4476, AB_2183326 at 1:500). Alexa Fluor-labeled secondary antibodies were used in parallel with Alexa Fluor 488 phalloidin (Invitrogen, A12379) to label F-actin. The secondary antibodies used were donkey anti-mouse Alexa Fluor 555 (Invitrogen, A32773), donkey anti-rat Alexa Fluor 488 (Invitrogen, A48269), donkey anti-rabbit Alexa Fluor 488 (Invitrogen, A32790), donkey anti-rabbit Alexa Fluor 555 (Invitrogen, A32794), donkey anti-goat Alexa Fluor 555 (Invitrogen, A32816), donkey anti-goat Alexa Fluor 647 (Invitrogen, A32849), and DAPI and Hoechst Nucleic Acid Stains (Invitrogen, D1306) for nuclear counterstain.

### Multiplex single molecule fluorescence *in situ* hybridization (smFISH)

Embryos were collected from timed pregnant mice at specified embryonic time points. Inner ears were dissected from embryos and fixed overnight at 4°C in 4% PFA in PBS. Inner ears for sectioning were equilibrated in sucrose, mounted in OCT, and frozen prior to cryosectioning. Sections were cut at a thickness of 12 μm. Hybridization was carried out using the RNAscope® Fluorescent Multiplex Reagent Kit v2 in accordance with the manufacturer’s guidelines (ACDbio). For whole mount smFISH, inner ears were isolated, fixed overnight, dehydrated through a graded MeOH/PBT series (25%, 50%, 75%, 100%), and stored at −20 °C. Inner ears were rehydrated through a reverse gradient of MeOH/PBT, followed by whole-mount smFISH as described in Kersigo et al.^87^ The probes we used were Mm-*Etv5* (ACDbio, #316961), Mm-*Etv1*-O1-C3 (ACDbio, #557891-C3), Mm-*Etv4*-C2 (ACDbio, #458121-C2), Mm-*Sox2*-C3 (ACDbio, #401041-C3), Mm-*Sox2*-C2 (ACDbio, #401041-C2), Mm-*Prox*-C2 (ACDbio, #488591-C2), Mm-*Prox1*(ACDbio, #401041), Mm-*Atoh1*-C2 (ACDbio, #408791-C2), Mm-*Mdm1*-C1 (ACDbio, #1784631-C1), Mm-*Sp5* (ACDbio, #405681), Mm-*Npnt* (ACDbio, #316771), Mm-*Lypd6* (ACDbio, #894701).

## Resource Availability

Please contact Matthew Kelley, with any requests

## Data and Code Availability

The original dataset described in this study has been deposited on GEO, accession number: GSE292561. In addition, the data will also be made available through gEAR (www.umgear.org). Other source data will be made available through Dryad.

## Acknowledgments

The authors would like to acknowledge the outstanding animal care provided by the staff of the PNRC Shared Animal Facility and the exceptional assistance of the members of the Genomics and Computational Biology Core at the NIDCD (ZIC DC 000086). This research was supported in part by the Intramural Research Program (ZIC DC000059 to MWK) of the National Institutes of Health (NIH). The contributions of the NIH author(s) are considered Works of the United States Government. The findings and conclusions presented in this paper are those of the author(s) and do not necessarily reflect the views of the NIH or the U.S. Department of Health and Human Services and by a grant from the Japanese Society for the Promotion of Science to SS. This study utilized the high-performance computational capabilities of the Biowulf Linux cluster at the National Institutes of Health, Bethesda, MD. (http://biowulf.nih.gov). The authors want to thank Emma R. Andersson, Doris Wu, Elizabeth Driver and Tessa Sanders for reading an earlier version of this manuscript and Braulio Peguero for advice on statistical analysis.

## Author Contributions

SS and MWK designed the study, SS conducted the experiments, analyzed the results, and performed bioinformatic analysis, SS and MWK prepared figures and wrote the manuscript.

