## Supplemental data for "Single-Cell Profiling of the Developing Organ of Corti Identifies Etv4/5/1 as Key Regulators of Pillar Cell Identity"

### Sakamoto and Kelley, Supplemental Figure 1

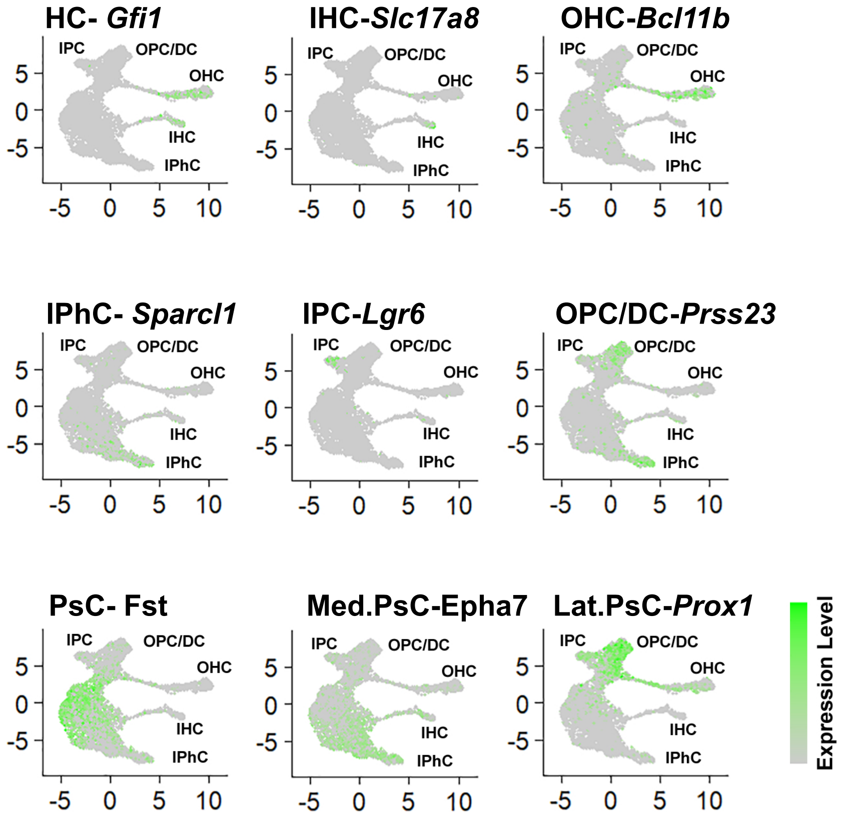

Sakamoto and Kelley, Supplemental Figure 2

#### PsC to Lat.PsC Transition

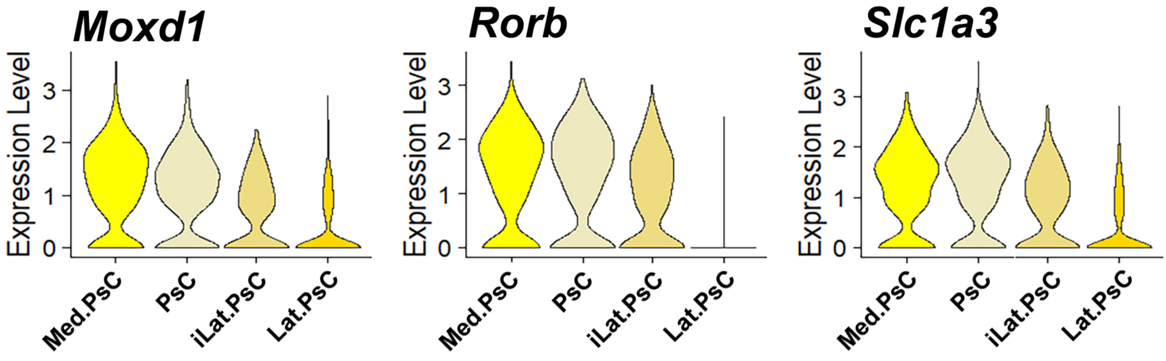


#### PsC to lPhC Transition

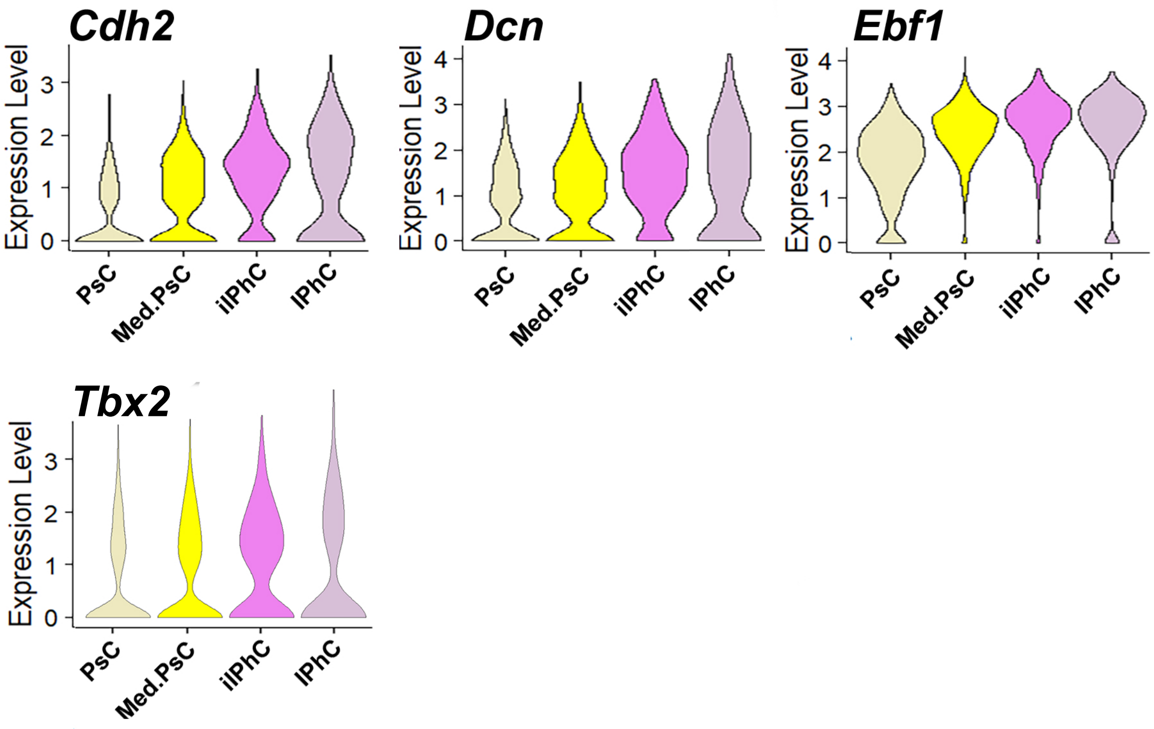

### Sakamoto and Kelley, Supplemental Figure 3

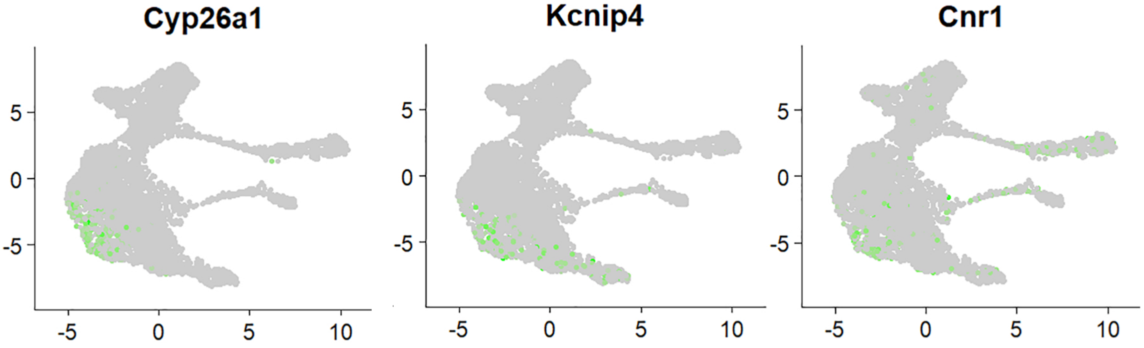

**Sakamoto and Kelley, Supplemental Figure 4**

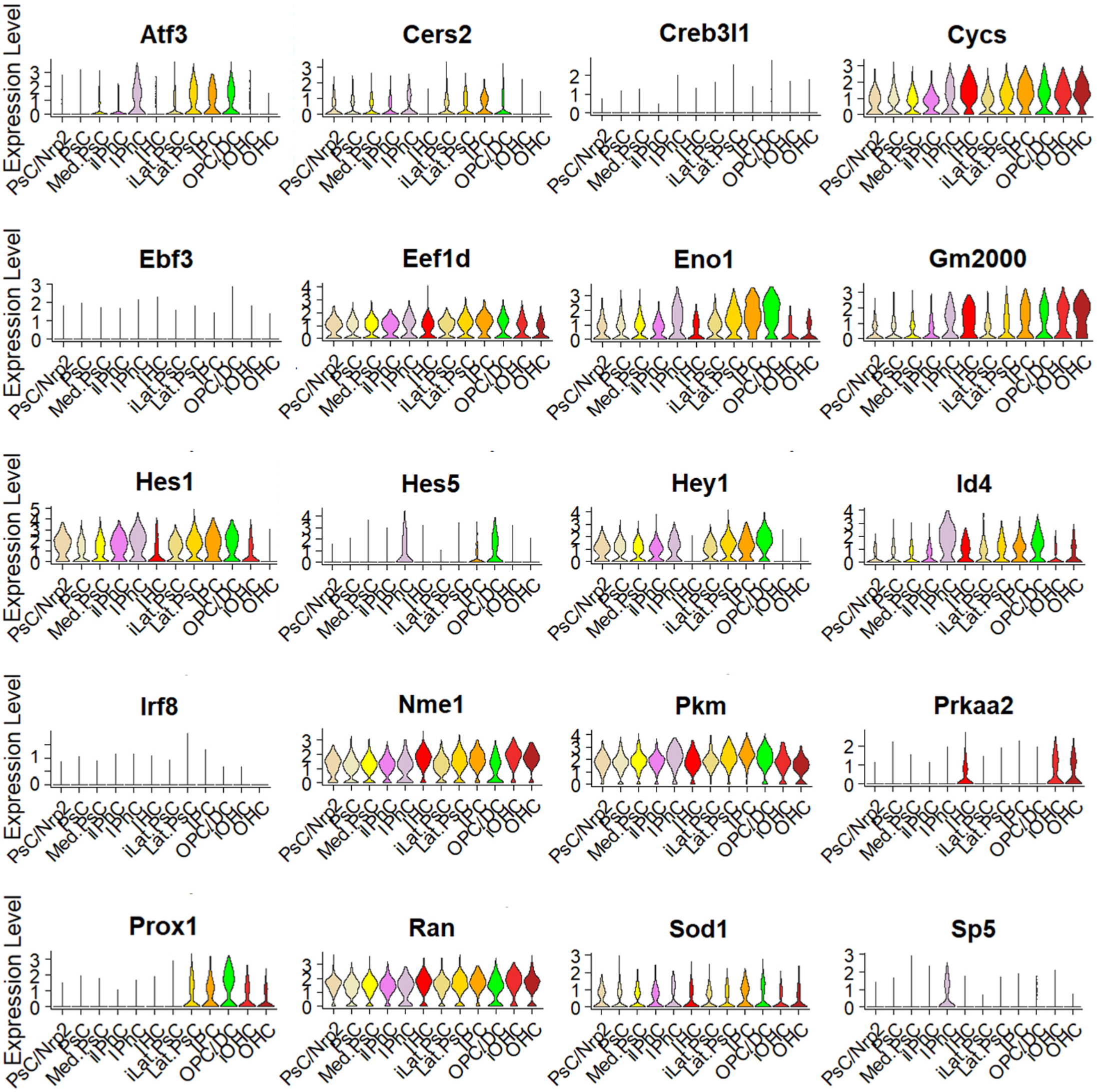

**Sakamoto and Kelley, Supplemental Figure 5**

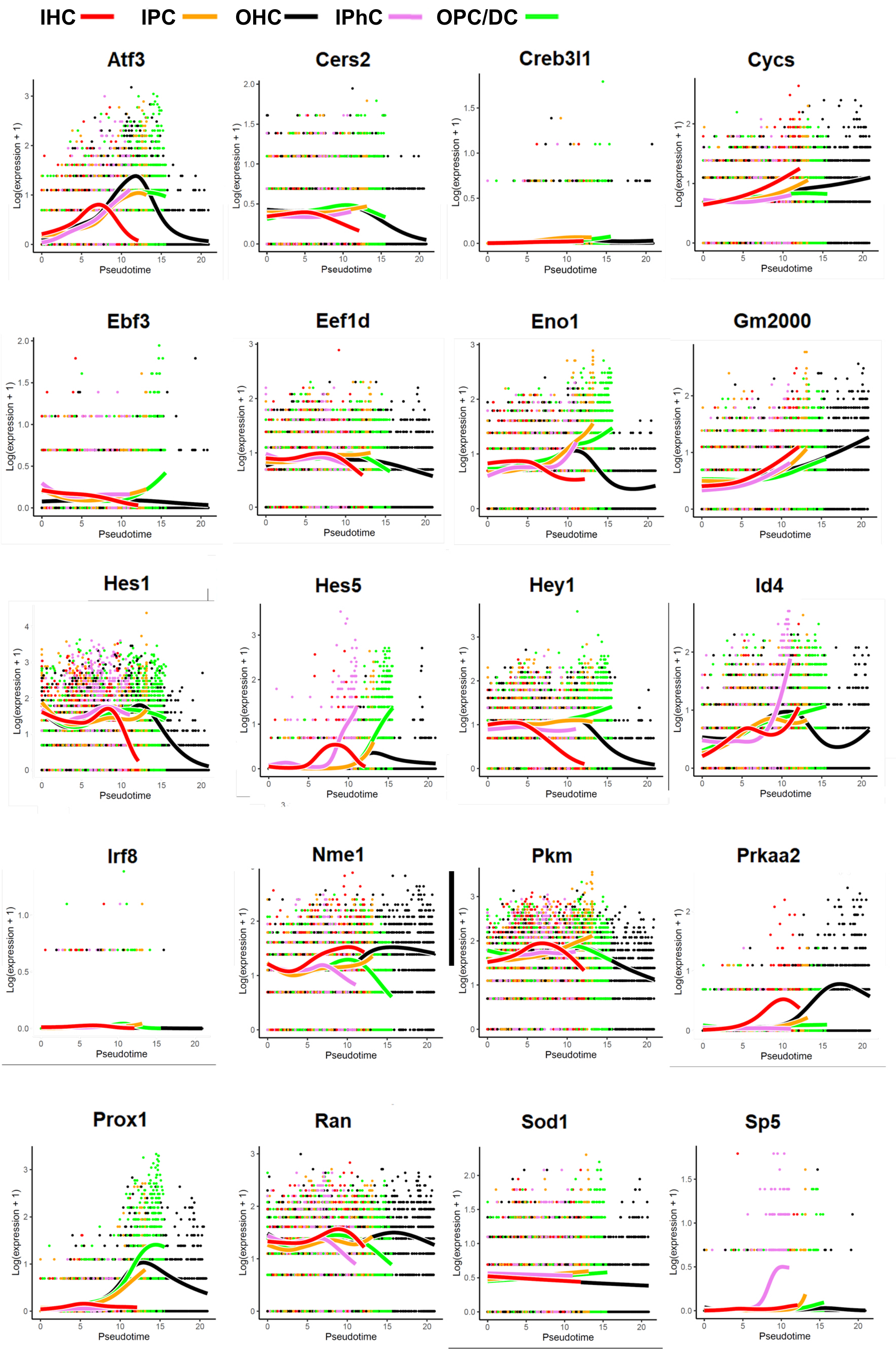

### Sakamoto and Kelley, Supplemental Figure 6

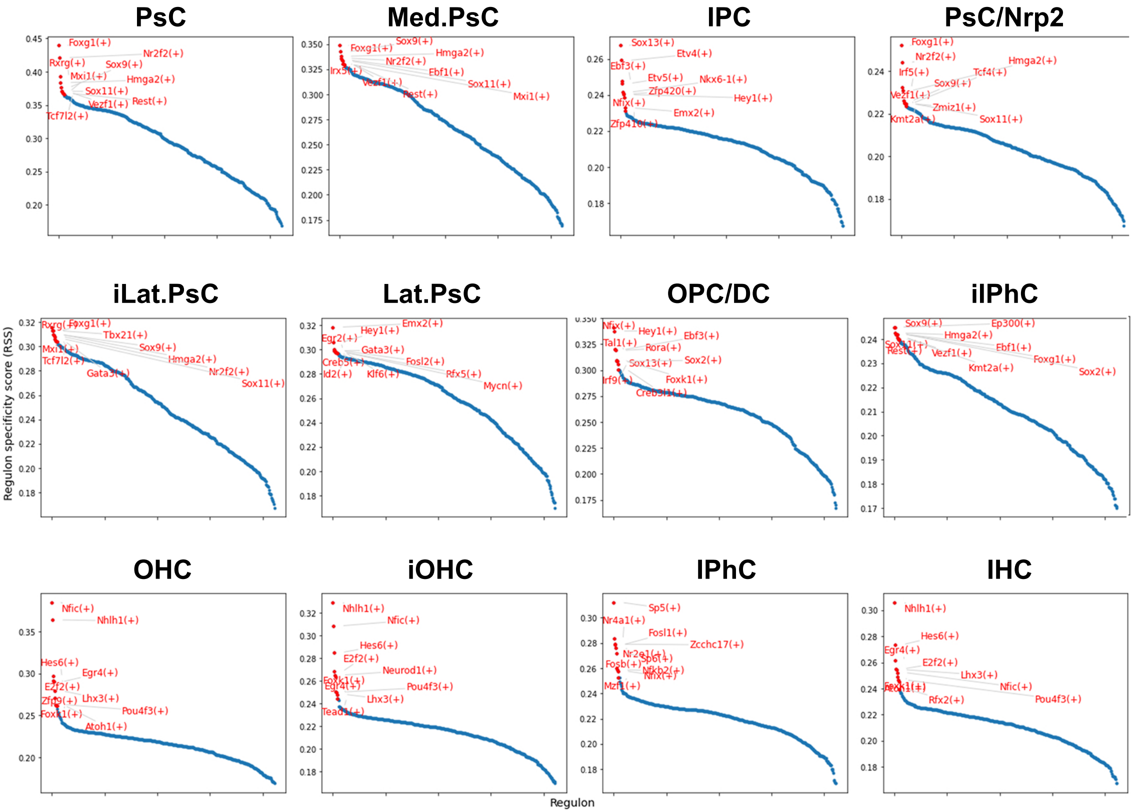

Sakamoto and Kelley, Suppl. Fig. 7

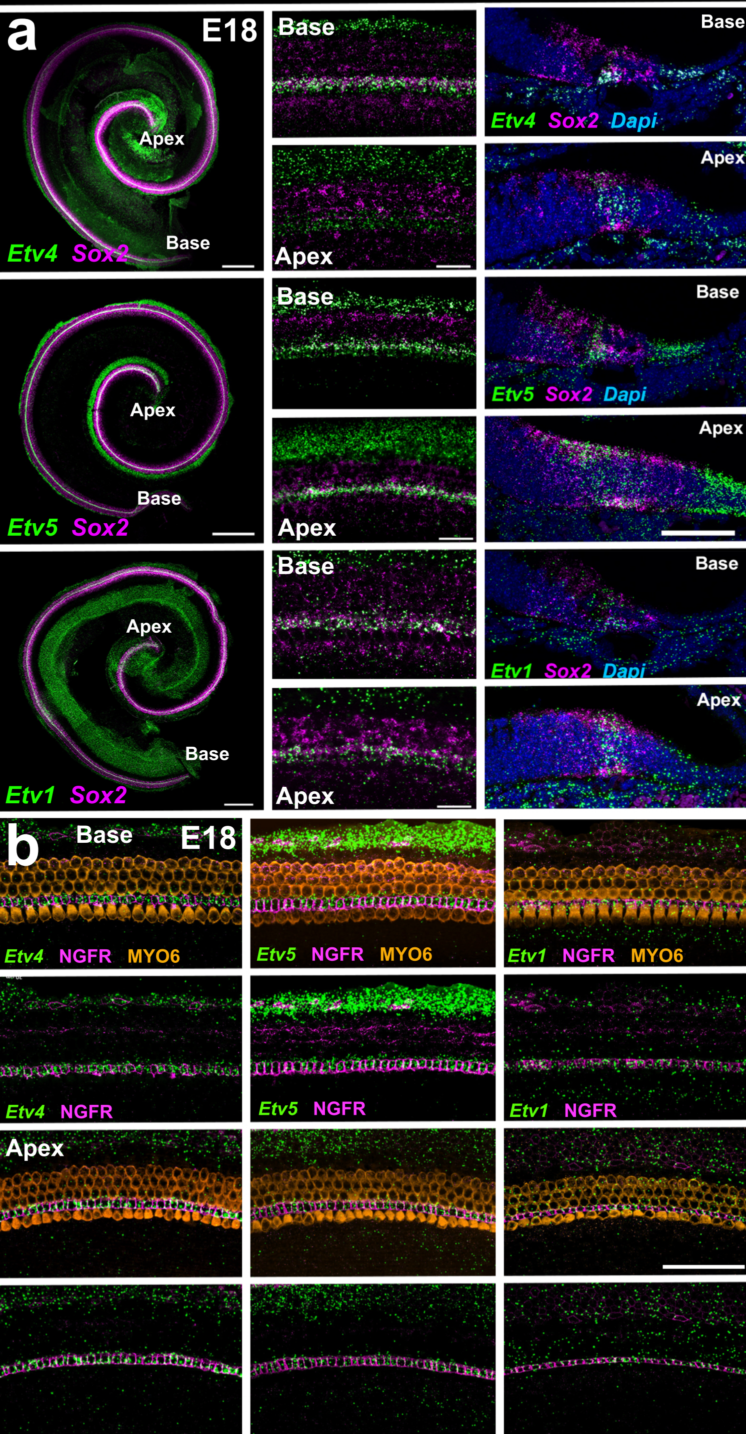

Sakamoto and Kelley, Supplemental Figure 8

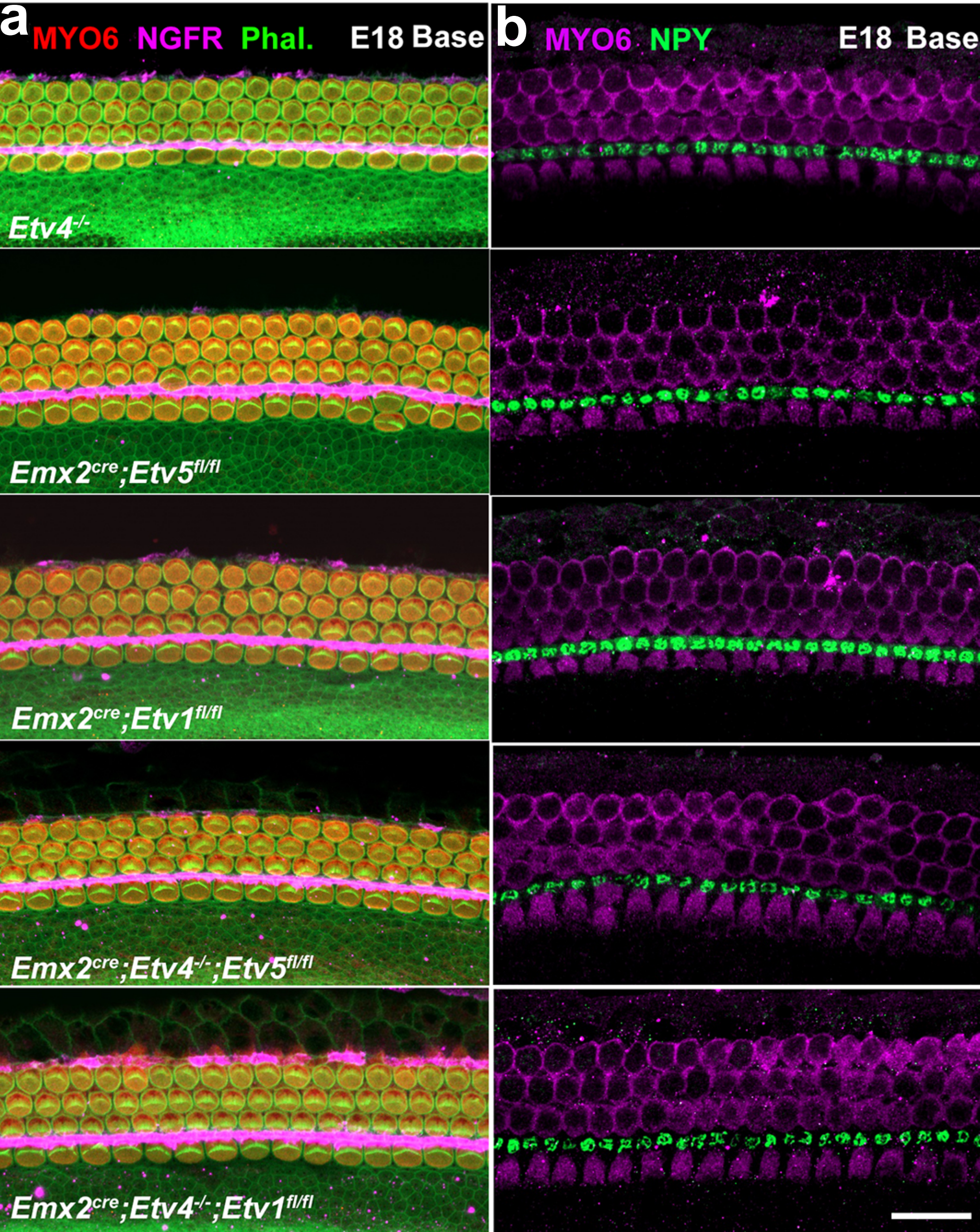

Sakamoto and Kelley, Supplemental Figure 9

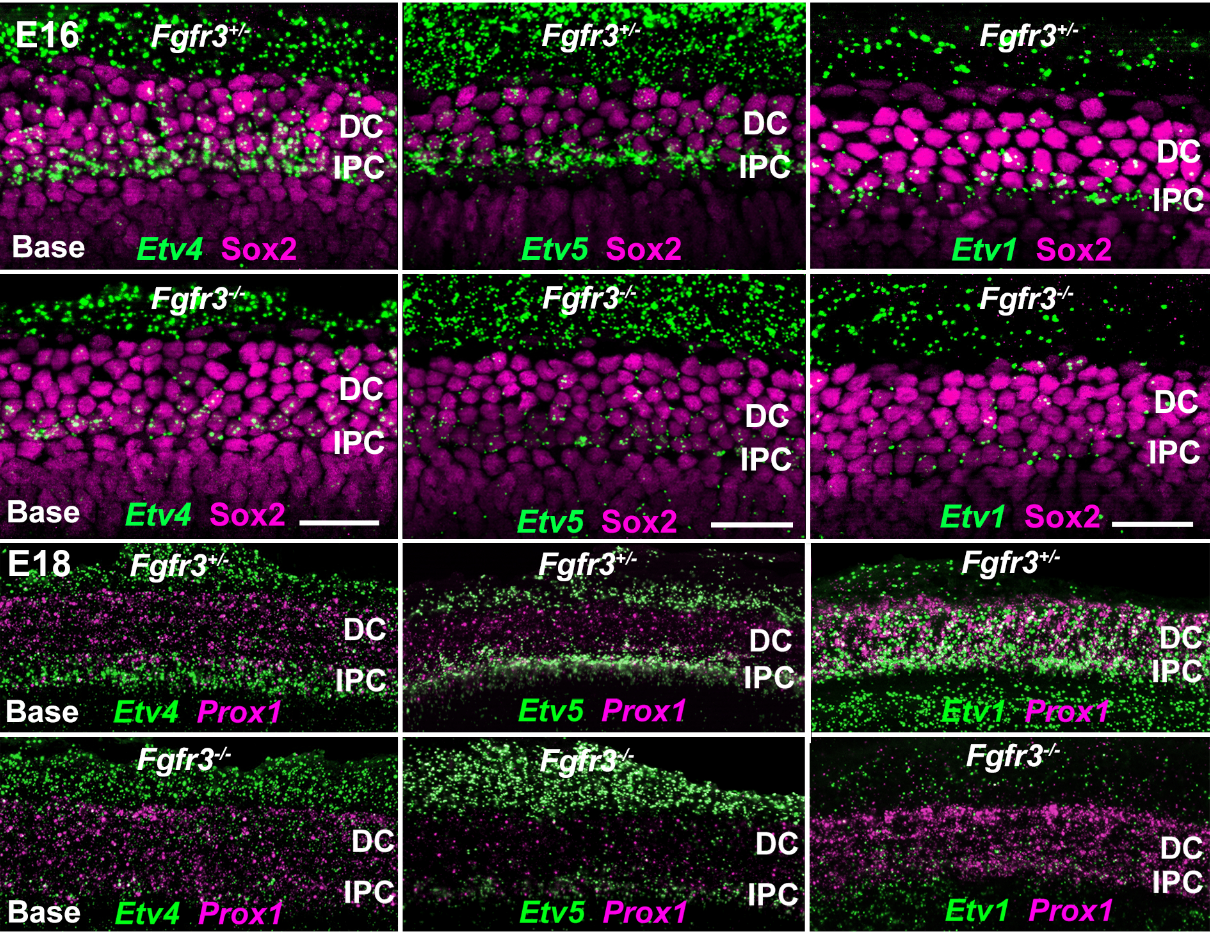

### Sakamoto and Kelley, Supplemental Figure 10. Changes in *Etv* Target Gene Expression in *Fgfr3*$^{-/-}$ Cochleae

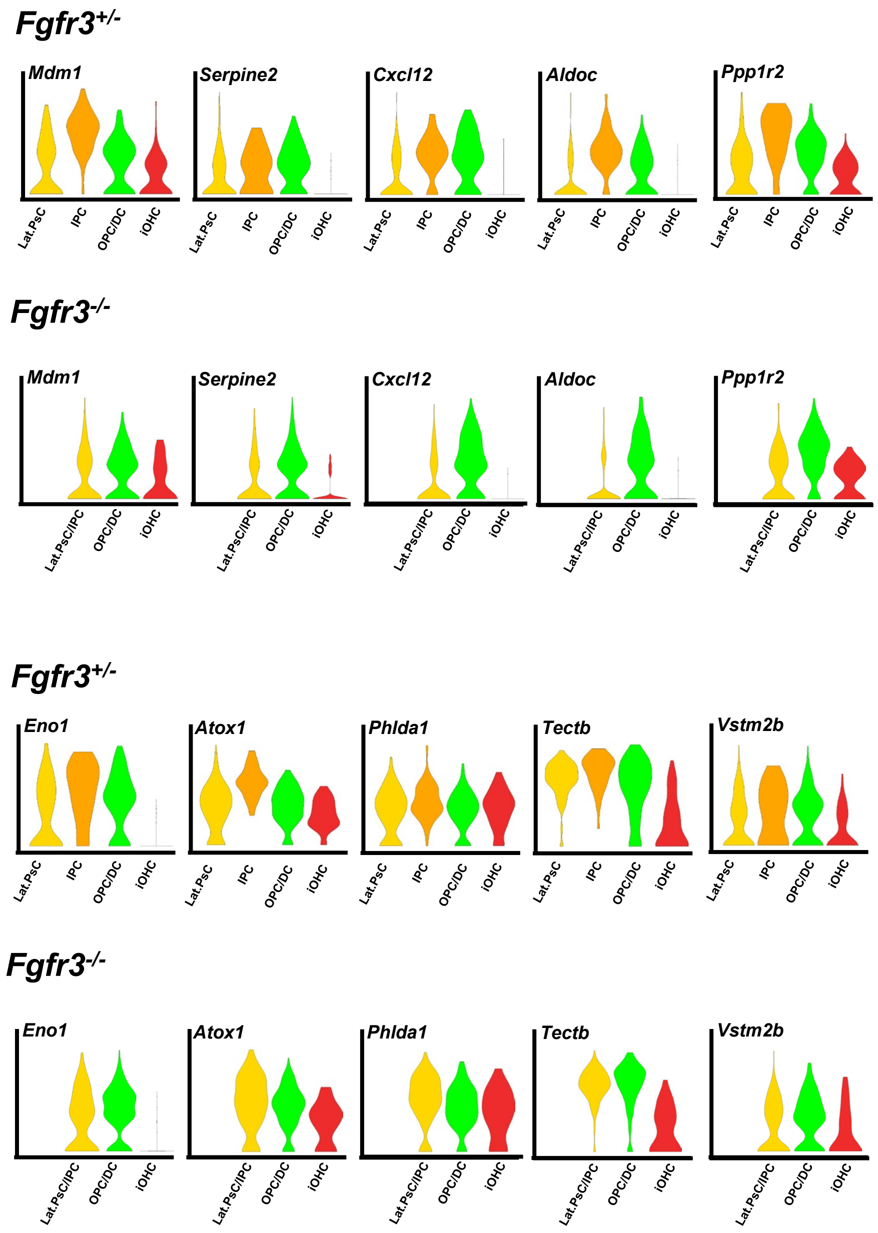

### Sakamoto and Kelley, Supplemental Figure 11.
### Changes in OHC and OPC/DC Gene Expression in *Fgfr3*$^{-/-}$ Cochleae

#### OHC Genes

##### *Fgfr3*$^{+/-}$

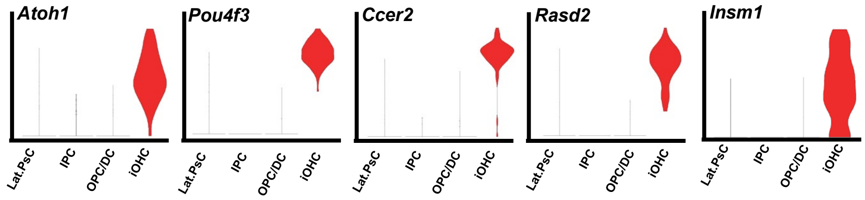


##### *Fgfr3*$^{-/-}$

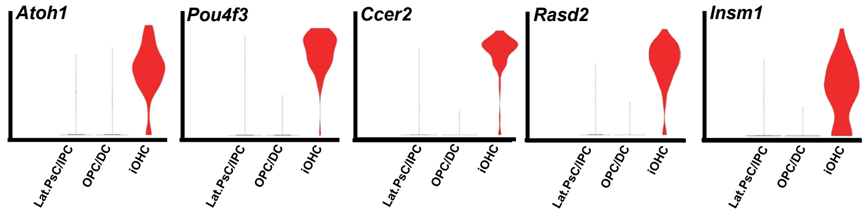


#### OPC/DC Genes

##### *Fgfr3*$^{+/-}$

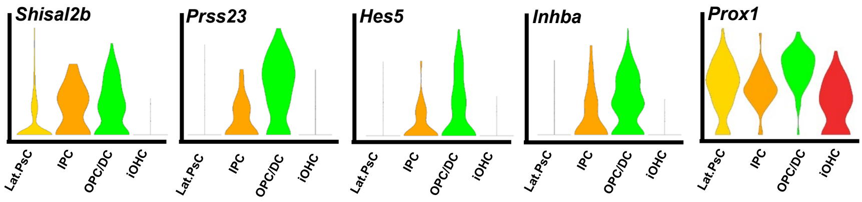


##### *Fgfr3*$^{-/-}$

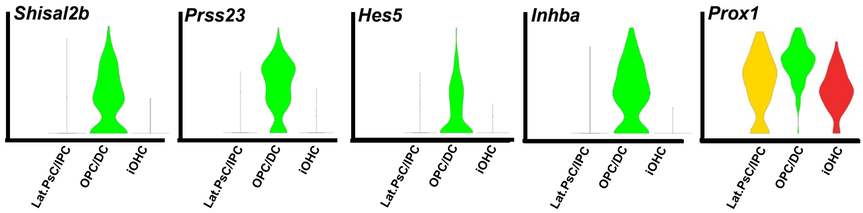
